# Cooperativity, information gain, and energy cost during early LTP in dendritic spines

**DOI:** 10.1101/2023.10.03.560651

**Authors:** Jan Karbowski, Paulina Urban

## Abstract

We investigate a mutual relationship between information and energy during early phase of LTP induction and maintenance in a large-scale system of mutually coupled dendritic spines with discrete internal states and probabilistic dynamics within the framework of nonequilibrium stochastic thermodynamics. In order to analyze this computationally intractable stochastic multidimensional system, we introduce a pair approximation, which allows us to reduce the spine dynamics into a lower dimensional manageable system of closed equations. It is found that the rates of information gain and energy attain their maximal values during an initial period of LTP (i.e. during stimulation), and after that they recover to their baseline low values, as opposed to a memory trace that lasts much longer. This suggests that learning phase is much more energy demanding than the memory phase. We show that positive correlations between neighboring spines increase both a duration of memory trace and energy cost during LTP, but the memory time per invested energy increases dramatically for very strong positive synaptic cooperativity, suggesting a beneficial role of synaptic clustering on memory duration. In contrast, information gain after LTP is the largest for negative correlations, and energy efficiency of that information generally declines with increasing synaptic cooperativity. We also find that dendritic spines can use sparse representations for encoding of long-term information, as both energetic and structural efficiencies of retained information and its lifetime exhibit maxima for low fractions of stimulated synapses during LTP. Moreover, we find that such efficiencies drop significantly with increasing the number of spines. In general, our stochastic thermodynamics approach provides a unifying framework for studying, from first principles, information encoding and its energy cost during learning and memory in stochastic systems of interacting synapses.

## 1. Introduction

Parts of synapses known as dendritic spines play an important role in learning and memory in neural networks (Bonhoeffer and Yuste 2002; Kasai et al 2003; Bourne and Harris 2008; Takeuchi et al 2014; Kandel et al 2014). Learning can be thought as acquiring information in synapses through plasticity mechanism such as LTP and LTD (long term potentiation and depression, respectively), and memory can be regarded as storing that information (for experimental overview see e.g.: Yang et al 2009; Bourne and Harris 2008; Takeuchi et al 2014; Poo et al 2016; for theoretical overview see e.g.: Fusi et al 2005; Benna and Fusi 2016; Chaudhuri and Fiete 2016). Thus learning and memory are strictly related to processing and maintaining of long term information, which in principle could be quantified in terms of information theory and statistical mechanics, similar as it was done for neural spiking activity (Rieke et al 1999). Indeed, recent results by the authors show that information (entropy) contained in the distributions of dendritic spine volumes and areas is nearly maximal for any given of their average sizes across different brains and cerebral regions (Karbowski and Urban 2022). This suggests that the concept of information can be useful in quantifying synaptic learning and memory, and that actual synapses might “use” and optimize certain information-theoretic quantities.

Physics teaches us that there are close relationships between information and thermodynamics, and the smaller the system the stronger the mutual link, since smallness enhances fluctuations and thus unpredictability in the system (Bennett 1982; Leff and Rex 1990; Berut et al 2012; Parrondo et al 2015). This means that information processing always requires some energy, and it is reasonable to assume that biological evolution favors systems that save energy while handling information, because of the limited resources and/or competition (Niven and Laughlin 2008). This line of thought was explored in neuroscience for estimating energy-efficient coding capacity in (short-term) neural activities and synaptic transmissions (Rieke et al 1999; Levy and Baxter 1996 and 2002; Levy and Calvert 2021; Laughlin et al 1998; Balasubramanian et al 2001). All these approaches and calculations for neural and synaptic activities, however valuable, missed one key ingredient of real biological systems, or did not make it explicit. Namely, biological systems, including neurons and synapses, always operate far from thermodynamic equilibrium, where balance between incoming and outgoing energy and matter (or probability) fluxes is broken; the so-called broken detailed balance (for a general physical approach, see: Maes et al 2000; Seiferet 2012; Gardiner 2004; for a biophysical approach to synapses, see: Karbowski 2019). As a consequence, equilibrium thermodynamics (with static or stationary variables and Gibbs distributions) does not seem to be the right approach, and to be more realistic, one has to use nonequilibrium statistical mechanics, where the concepts of stochasticity and entropy production play central roles (and where we do not know a priori the probability distributions). Nonequilibrium statistical mechanics (or nonequilibrium stochastic thermodynamics) provides a unifying description for stochastic dynamics, because it treats microscopic information and energy on the same footing, which allows us to get the right estimates of information and energy rates from first principles. Such an approach was initiated in (Karbowski 2019, 2021) for studying nonequilibrium thermodynamics of synaptic plasticity. Both of these studies suggested that synaptic plasticity can use energy rather economically, since (i) it consumes only about 4 − 11% of energy devoted for fast synaptic transmission (Karbowski 2019), and (ii) it can provide higher coding accuracy and longer memory time for a lower energy cost at certain regimes (Karbowski 2019, 2021).

There exists a large experimental evidence that local synaptic cooperativity on a dendrite takes place during learning and memory formation (Marino and Malinow 2011; Yadav et al 2012; for a review see: Winnubst et al 2012), and it can also be useful for long-term memory stability (Govindarajan et al 2006; Kastellakis and Poirazi 2019). Thus, it seems that any realistic model of synaptic plasticity relevant for memory formation should include correlations between neighboring synapses. Moreover, it would be good to know how such correlations influence the lifetime of memory trace, as well as, information gain and energy cost associated with it.

The current study explores nonequilibrium statistical mechanics for investigating the efficiency of learning and storing information in a system of interacting synapses with stochastic dynamics during early LTP induction and its maintenance (e-LTP phase, without consolidation). Not only fast synaptic transmission is noisy (Volgushev et al 2004); the noise is also present in long-term synaptic dynamics associated with slow plasticity due to large thermal fluctuations in internal molecules and presynaptic input (Bonhoeffer and Yuste 2002; Holtmaat et al 2005; Choquet and Triller 2013; Statman et al 2014; Meyer et al 2014; Kasai et al 2003; Loewenstein et al 2011). Thus synaptic plasticity requires a probabilistic approach (Yasumatsu et al 2008), and we assume that it can be described as dynamics involving transitions between discrete mesoscopic states (e.g. Montgomery and Madison 2004; Fusi et al 2005; Leibold and Kempter 2008; Barrett et al 2009; Benna and Fusi 2016). Our paper extends the previous two studies (Karbowski 2019, 2021) in three important methodological ways. First, it provides a general framework for approximating the dynamics of multidimensional stochastic system of *N* interacting synapses each with 4 internal states (which in practice is computationally intractable for large *N*), by reducing it to a set of coupled lower-dimensional stochastic subsystems (which are computationally tractable). This is done by applying the so-called pair approximation, which allows us to reduce the system with 4^*N*^ equations to the system with ∼ 4(5*N* − 4) equations. Second, our paper provides explicit formulas for investigating information gain (rate of Kullback-Leibler divergence) and energy consumption (entropy production rate) for arbitrary time during learning and memory retention phases. Third, we use data-driven estimates for transition rates between different synaptic states, and hence our values of information and energy are realistic.

On a conceptual level, our study investigates how the efficiency of encoded information (both amount and duration) depends on a degree of correlation between neighboring synapses, on a percentage of their activation by presynaptic neurons, and on magnitude and duration of synaptic stimulation during LTP. The first relates to cooperativity between synapses, the second to sparseness of synaptic coding, and the last to the strength of learning. All these parameters can in principle be compared to empirical data, once those are available, thus providing an important link between theory and experiment.

Throughout the paper we use interchangeably the terms “dendritic spine” and “synapse”.

## 2. Model of plastic interacting dendritic spines

We consider a single postsynaptic neuron having one basal (main) dendrite with *N* dendritic spines located along its length (Fig. 1). Because of the small sizes of dendritic spines (∼ 1*μ*) their dynamics is necessarily probabilistic due to thermodynamic fluctuations of local environment (with high temperature ∼ 300 K), as well as due to activity fluctuations in neurons (both of electric and chemical nature). Moreover and importantly, the spines are locally coupled by nearest neighbor interactions, as the experimental data suggest (Marino and Malinow 2011; Yadav et al 2012; Winnubst et al 2012)).

**Fig. 1.**
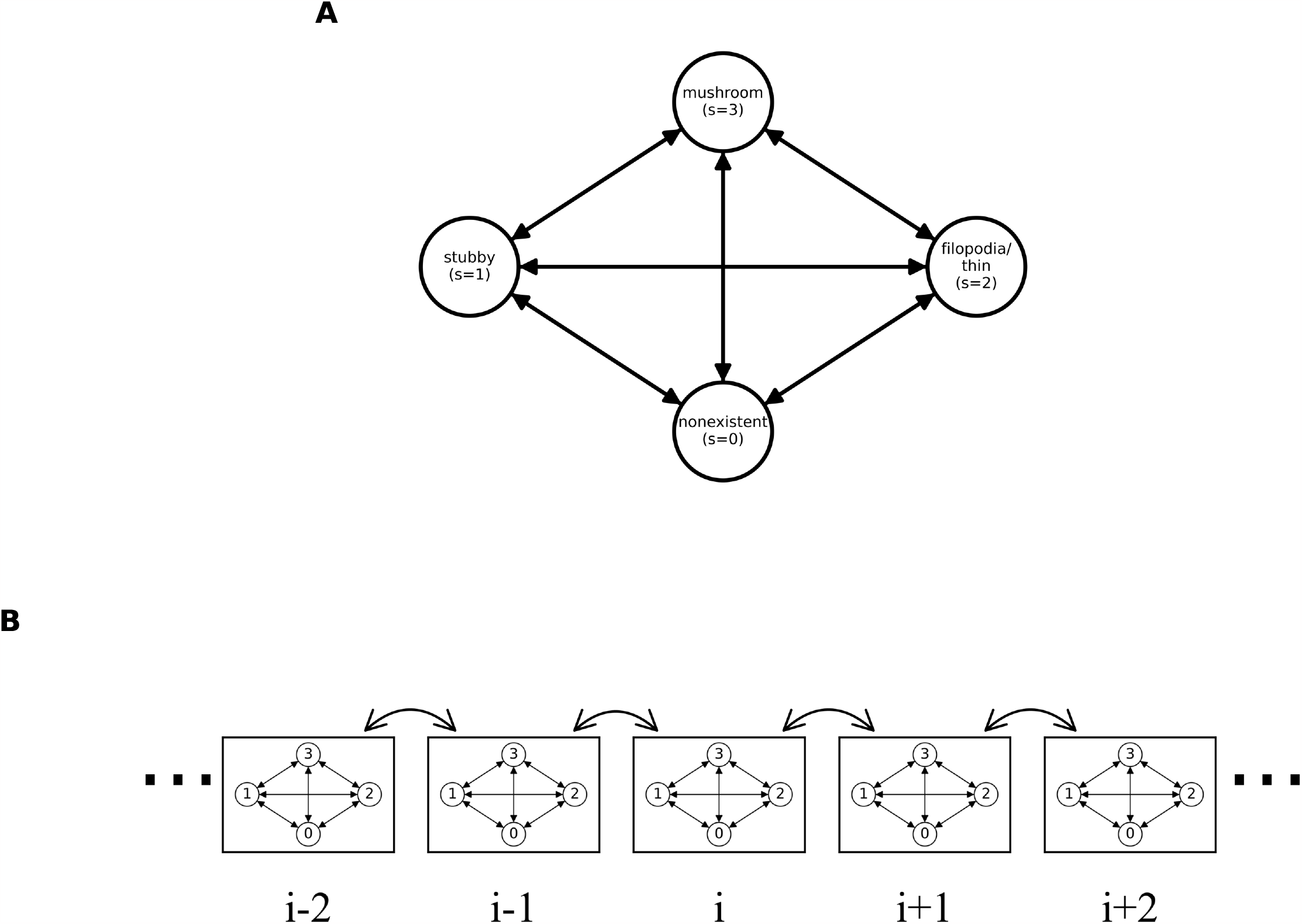
Mesoscopic model of dendritic spines and their interactions on a dendrite. A) Four state stochastic model of a dendritic spine with transitions between the mesoscopic states. B) Interactions between spines on a dendrite are determined only by nearest neighbors, in such a way that spine’s intrinsic transition rates depend on the states of neighboring spines. These interactions can be thought as representing inflow and outflow of different molecules between spines.

### 2.1 Morphological spine states and stochastic multidimensional dynamics

Empirical data indicate that a single dendritic spine can be regarded as a four state stochastic system with well defined morphological (mesoscopic) states (Montgomery and Madison 2004; Bokota et al 2016; Basu et al 2018; Urban et al 2020; Fig. 1A). These states are denoted here as *s*_*i*_ at each location *i*, with values *s*_*i*_ = 0, 1, 2, 3, corresponding respectively to the following morphological states: nonexistent, stubby, filopodia/thin, and mushroom (the larger *s*_*i*_, the larger the spine size; see Appendix A). These mesoscopic states are quasi stable, which means that there are slow transitions between them that are much slower than microscopic transitions between molecular, mostly unknown, processes comprising internal microscopic dynamics of a dendritic spine (Kennedy 2000; Sheng and Hoogenraad 2007; Miller et al 2005; Kandel et al 2014). This approach can be treated as a coarsegrained description of intrinsic spine dynamics in terms of stochastic Markov process on a mesoscopic scale.

We assume stochastic dynamics for *N* coupled dendritic spines, and denote by *P* (*s*_1_, *s*_2_, …, *s*_*N*_ ; *t*) the probability that the spine system has the configuration of internal states *s*_1_, *s*_2_,, *s*_*N*_ . The dynamic of this global stochastic state is motivated by the Glauber model of time-dependent Ising model (Glauber 1963), and it is represented by the following Master equation (see Appendix A)

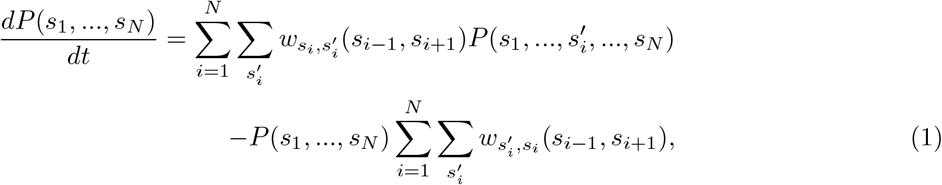

where 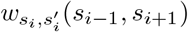 is the transition rate for the jumps inside spine *i* from state 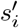 to state *s*_*i*_.These jumps also depend on the states of neighboring spines *s*_*i*−1_ and *s*_*i*+1_, because of the nearest neighbor coupling between the spines (Fig. 1B). Generally, we take the following form of the transition matrix 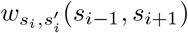:

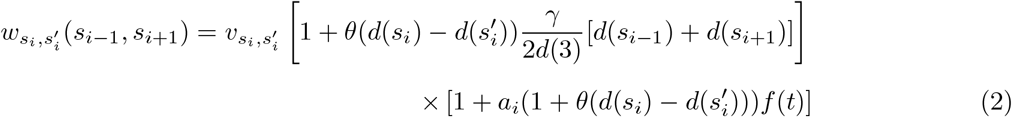

where 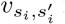 is the intrinsic basic transition rate between 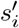 and *s*_*i*_ at spine *i*, and it is independent of the neighboring synapses. These intrinsic rates are the same for each *i*, setting the temporal scale for basal synaptic plasticity, and they were estimated based on data in (Urban et al 2020) and are presented in Table 1. The symbol *d*(*s*_*i*_) is the spine size at state *s*_*i*_ of spine *i* and it is proportional to spine surface area (see Appendix A), whereas *θ*(*x*) denotes the sign function of the argument *x*, where *θ*(*x*) = 1 if *x* ≥ 0, and *θ*(*x*) = −1 if *x <* 0. Note that larger spine sizes of neighboring spines generally influence more the transition rates of the given spine, which relates to their cooperativity. The parameter *γ* corresponds to magnitude of spine cooperativity between nearest neighbors, with −1 ≤ *γ* ≤ 1, and there is rescaling by the maximal spine size *d*(3) (size in state *s* = 3 called mushroom), which ensures positivity of all elements of the transition matrix. The positive values of *γ* indicate positive correlations (positive cooperativity), while its negative values mean negative correlations (negative cooperativity). Note that for positive cooperativity, the local spine interactions amplify the transitions that lead to increase of spine size, and reduce those transitions that decrease spine size. The opposite is true for negative cooperativity. It should be added that distant spines also can affect a given spine at location *i*, but that interaction has an indirect character and thus is weaker and mediated with some delay.

**Table 1:** Values of intrinsic basic transition rates *v*_*s,s*_′ in each synapse.

| Matrix element | Value |
| --- | --- |
| $v_{0,1}$ | 0.019 |
| $v_{1,0}$ | 0.065 |
| $v_{0,2}$ | 0.015 |
| $v_{2,0}$ | 0.007 |
| $v_{0,3}$ | 0.022 |
| $v_{3,0}$ | 0.001 |
| $v_{2,1}$ | 0.003 |
| $v_{1,2}$ | 0.061 |
| $v_{2,3}$ | 0.009 |
| $v_{3,2}$ | 0.015 |
| $v_{1,3}$ | 0.049 |
| $v_{3,1}$ | 0.008 |

All diagonal elements, i.e. $v_{k,k}$ , are zero. The units are in $\text{min}^{-1}$ .

The last factor on the right in Eq. (2) indicates the effect of external (presynaptic) stimulation, leading to LTP, with a time varying function known as alpha function *f* (*t*) given by

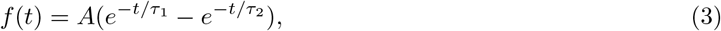

where *A* is the stimulation amplitude, *t* is the time after stimulation onset, and *τ*_1_, *τ*_2_ are time constants related to falling and rising phases of the stimulation, respectively. The latter means that LTP related stimulation last only about *τ*_1_ + *τ*_2_ (∼ 17 min; Table 2), which we call the duration of the learning phase. After that time, all the transition rates essentially decay to their pre-stimulation basal values. Consequently, after the stimulation, the dynamics of synaptic plasticity is driven only by the interactions between neighboring spines and their internal states. These dynamics are slow, and we call this stage the memory phase. The prefactor *a*_*i*_ of *f* (*t*) in Eq. (2) assumes two values: *a*_*i*_ = 1 when the spine *i* is stimulated with the probability *p*_*act*_, and *a*_*i*_ = 0 when there is no stimulation with the probability 1 − *p*_*act*_. Note that LTP related stimulation amplifies only the transitions increasing the spine size (the sign function *θ* is then 1). For the transitions decreasing the spine size, there is no amplification, because then the prefactor of *f* (*t*) is zero.

**Table 2:** Nominal values of global parameters used in computations.

| Description | Variable | Value | Units |
| --- | --- | --- | --- |
| Number of synapses | $N$ | 1000 | unitless |
| Cooperativity | $\gamma$ | (-1.0,1.0) | unitless |
| Probability of stimulation | $p_{act}$ | 0.3 | unitless |
| Stimulation amplitude | $A$ | 200 | unitless |
| Decay time of stimulation | $\tau_1$ | 15 | min |
| Rising time of stimulation | $\tau_2$ | 2 | min |
| Energy scale for plasticity | $\epsilon$ | $4.6 \cdot 10^5$ | kT |

Value of stimulation amplitude $A$ is motivated by experimental facts that during LTP the rates of protein phosphorylation at PSD increase by 2 – 3 orders of magnitude in respect to baseline rates (e.g. Miller et al 2005). Energy scale $\epsilon$ for synaptic plasticity was taken from the estimate in Karbowski (2021).

The important point is that during learning (the phase when the function *f* (*t*) is activated), the information about the stimulation is encoded in the patterns of the probabilities *P* (*s*_1_, …, *s*_*N*_). Thus, knowing how these patterns change in time provides necessary “data” for computing physical characteristics of learning and memory.

### 2.2 Reduction of multidimensional spine stochastic dynamics into low dimensional dynamics: pair approximation

Equation (1) describes dynamics of the multidimensional probability that involves a gigantic 4^*N*^ number of equations. For a typical number of synapses on a dendrite *N* ∼ 10^3^, the dynamics represented by Eq. (1) require ∼ 10^600^ equations, which are impossible to simulate and solve on any existing computer. This numerical impossibility forces us to find an approximation to the dynamics in Eq. (1). Consequently, we consider a lower dimensional dynamics involving only singlets and pairs of locally interacting spines. This strategy is sufficient to describe the global dynamics of the synaptic system, and to compute information and energy rates, if we make a certain reasonable assumption (see below).

The probabilities for singlets and doublets of spine states *P* (*s*_*i*_) and *P* (*s*_*i*_, *s*_*i*+1_) can be obtained from Eq. (1) by summations over all other states in other synapses as

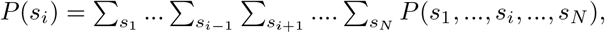

and similarly

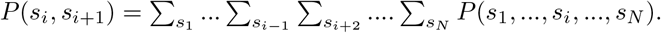

As a result we obtain two equations

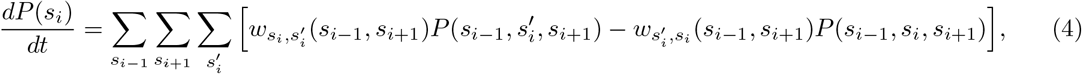

which is valid for *i* = 2, …, *N* − 1, and

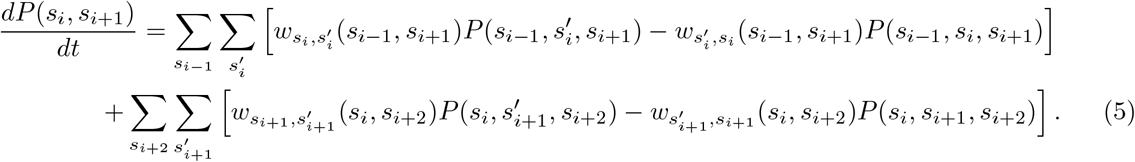

which is valid for *i* = 2, …, *N* − 2. For the boundary probabilities, i.e., for the dynamics of *P* (*s*_1_), *P* (*s*_*N*_) and *P* (*s*_1_, *s*_2_), *P* (*s*_*N*−1_, *s*_*N*_) we have similar equations, except we drop the boundary terms *s*_0_, *s*_*N*+1_ as they are nonexistent.

Equations (4) and (5) for the dynamics of *P* (*s*_*i*_) and *P* (*s*_*i*_, *s*_*i*+1_) involve additionally the probabilities of spine triplets *P* (*s*_*i*−1_, *s*_*i*_, *s*_*i*+1_) and *P* (*s*_*i*_, *s*_*i*+1_, *s*_*i*+2_), and thus they do not form a closed system of equations. To close these equations, we use the so-called “pair approximation” for probabilities. The main idea in this approximation is that the biggest influence on a given synapse is exerted only by the nearest neighbor synapses, and the effects from remote neighbors can be neglected, as is implied by the form of the transition rate 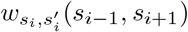. Specifically, for 3 neighboring synapses indexed spatially as *i* − 1, *i, i* + 1, the dynamic of synapse *i* − 1 depends directly only on the state of synapse *i*, and the influence of *i* + 1 synapse can be neglected as coming from the remote site. In terms of probabilities, this can be written as

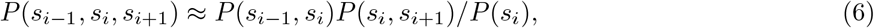

where we used the approximation for the conditional probability *P* (*s*_*i*−1_|*s*_*i*_, *s*_*i*+1_) ≈ *P* (*s*_*i*−1_|*s*_*i*_), and the fact that *P* (*s*_*i*−1_|*s*_*i*_) = *P* (*s*_*i*−1_, *s*_*i*_)*/P* (*s*_*i*_). Thus the probabilities of the spine triplets can be effectively written as combinations of the probabilities for spine singlets and doublets, which forms the essence of the pair approximation. A similar expression can be obtained for synapses with other combination of indexes.

The above pair approximation allows us to write the dynamics of probabilities *P* (*s*_*i*_) and *P* (*s*_*i*_, *s*_*i*+1_) as

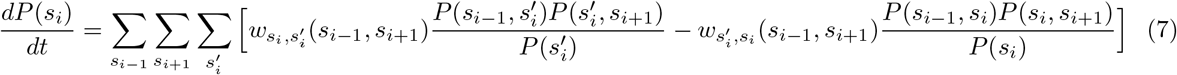

for *i* = 2, …, *N* − 1, and

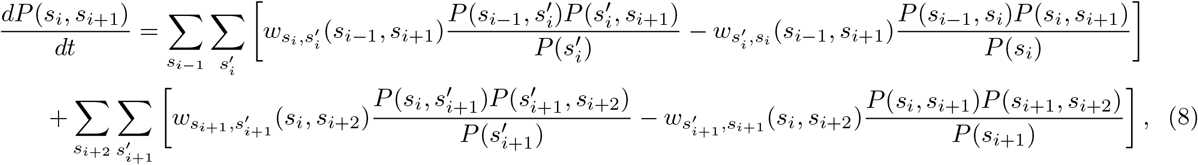

for *i* = 2, …, *N* − 2. Similar expressions can be written for the boundary probabilities with *i* = 1 and *i* = *N*.

It is clear that Eqs. (7,8) for the dynamics of *P* (*s*_*i*_) and *P* (*s*_*i*_, *s*_*i*+1_) form the closed system of equations, since now they only depend on each other. Importantly, the number of equations in the reduced dynamics (7,8) is only 20*N* − 16, which is linear in *N* and thus much smaller than the original 4^*N*^ equations, and hence feasible for numerical analysis. These two types of probabilities are sufficient to compute quantities of interest, which are associated with LTP induction, such as memory trace and its duration, the average sizes of spines, and the rates of information gain (Kullback-Leibler divergence) and energy dissipated (entropy production rate). However, first we check the accuracy of the pair approximation.

### 2.3 Validity of the pair approximation

In this section we check how accurate is the pair approximation by considering a small system of dendritic spines with *N* = 4, for which one can find an exact numerical solution for the dynamics in Eq. (1). Our goal is to compare this exact solution with its approximation given by Eqs. (7) and (8).

Numerical calculations indicate that the pair approximation is very accurate, as exact and approximate probabilities are practically indistinguishable (Fig. 2A), even for very strong couplings between spines (*γ* = −0.9 and *γ* = 0.9). Moreover, the pair approximation is well defined, since it preserves positivity of all probabilities and their normalization (Fig. 2A,B). Additionally, as an example of the main observable used in this study, we also compared entropy production rates computed for the exact dynamics in Eq. (1), denoted as EPR_*ex*_, with that computed from the approximate dynamics in Eqs. (7) and (8) and denoted as EPR_*pa*_ (the formulas for the exact and approximate EPR are given, respectively, in Eqs. (10-13) and Eqs. (16-19)). Both entropy production rates are also essentially indistinguishable, with a small difference at most 0.6% (Fig. 2C).

**Fig. 2.**
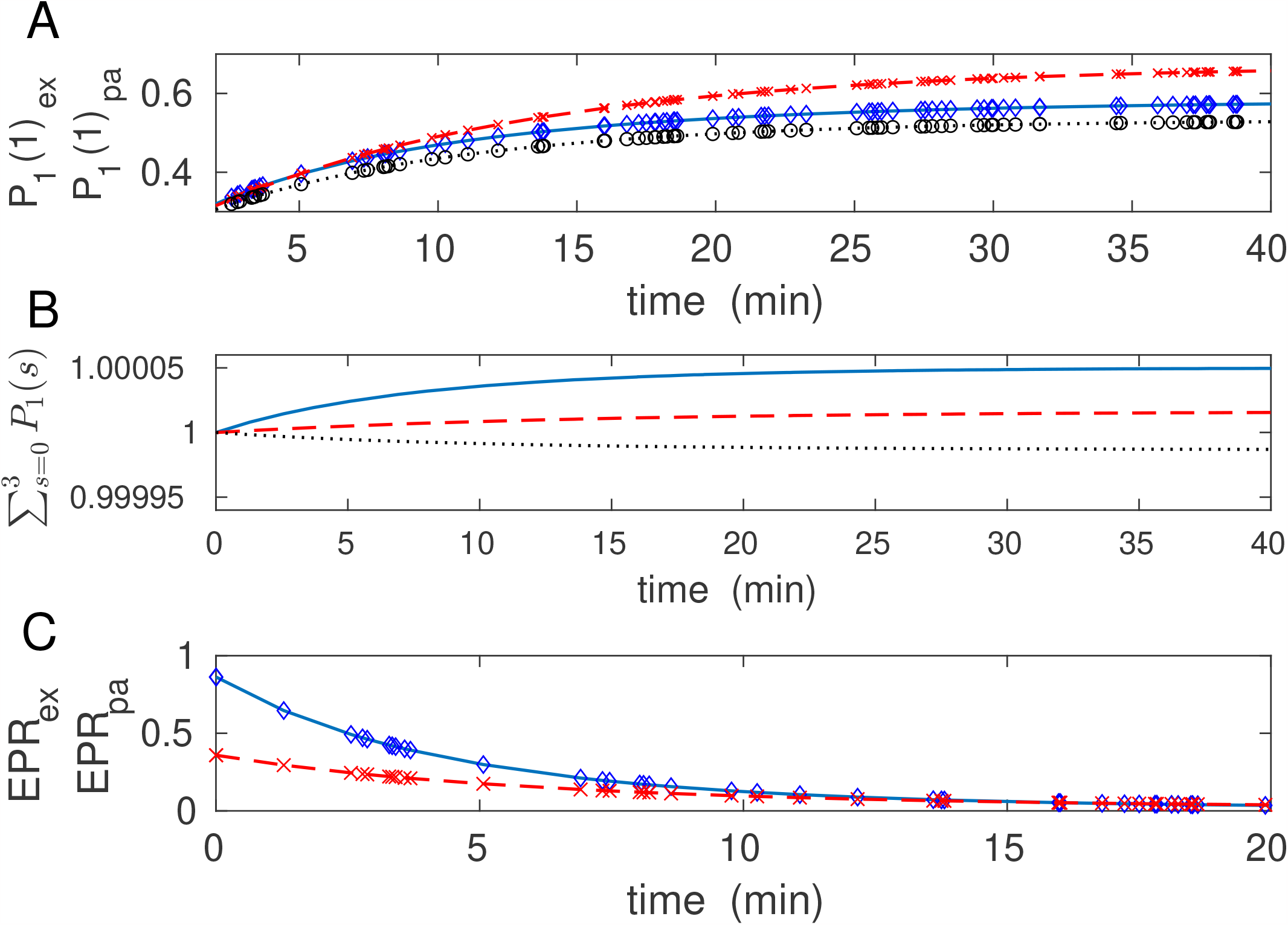
Comparison of an exact solution and pair approximation for N=4 interacting spines. A) Time dependence of exact *P*_1_(1)_*ex*_ and approximated *P*_1_(1)_*pa*_ probability *P* (*s*_1_ = 1) for three different couplings *γ*. Both probabilities are essentially indistinguishable. Exact solutions correspond to solid (*γ* = −0.9), dashed (*γ* = 0.1), and dotted (*γ* = 0.9) lines. Pair approximations correspond to diamonds (*γ* = −0.9), x (*γ* = 0.1), and circles (*γ* = 0.9). B) Normalization condition for probabilities of spine no 1, i.e. 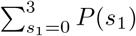. Solid line for *γ* = −0.9, dashed line for *γ* = 0.1, and dotted line for *γ* = 0.9.Note that the sum deviates from unity by a very small number less than 5 · 10^−5^. C) Time dependence of exact (EPR_*ex*_) and approximated (EPR_*pa*_) entropy production rate. Exact EPR_*ex*_ correspond to solid (*γ* = −0.9) and dashed (*γ* = 0.1) lines. Pair approximations EPR_*pa*_ correspond to diamonds (*γ* = −0.9) and x (*γ* = 0.1). Note an excellent matching of EPR_*pa*_ to EPR_*ex*_.

In order to give a measure of the pair approximation accuracy we introduce the ratio *R* for *N* = 4 spines, defined as

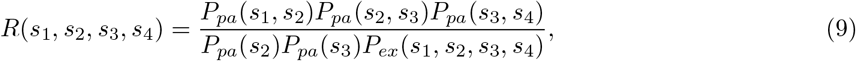

where the subscript *pa* refers to the pair approximation (Eqs. 7 and 8), while *ex* corresponds to the exact solution (Eq. 1). When *R* approaches 1, then the pair approximation matches the exact solution perfectly. The larger the deviation of *R* from unity, the less accurate is the approximation. This follows from the form of pair approximation for 4 spines, i.e., *P* (*s*_1_, *s*_2_, *s*_3_, *s*_4_) ≈ *P* (*s*_1_, *s*_2_)*P* (*s*_2_, *s*_3_)*P* (*s*_3_, *s*_4_)*/*[*P* (*s*_2_)*P* (*s*_3_)]. To have a global numerical accuracy, we have to average *R* over all states, which yields a mean ratio 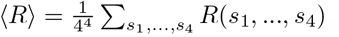, and its standard deviation 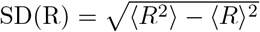, serving as a global error. In Fig. 3, we show that ⟨*R*⟩ is very close to 1, and SD(R) is generally small, at most 0.1.

**Fig. 3.**
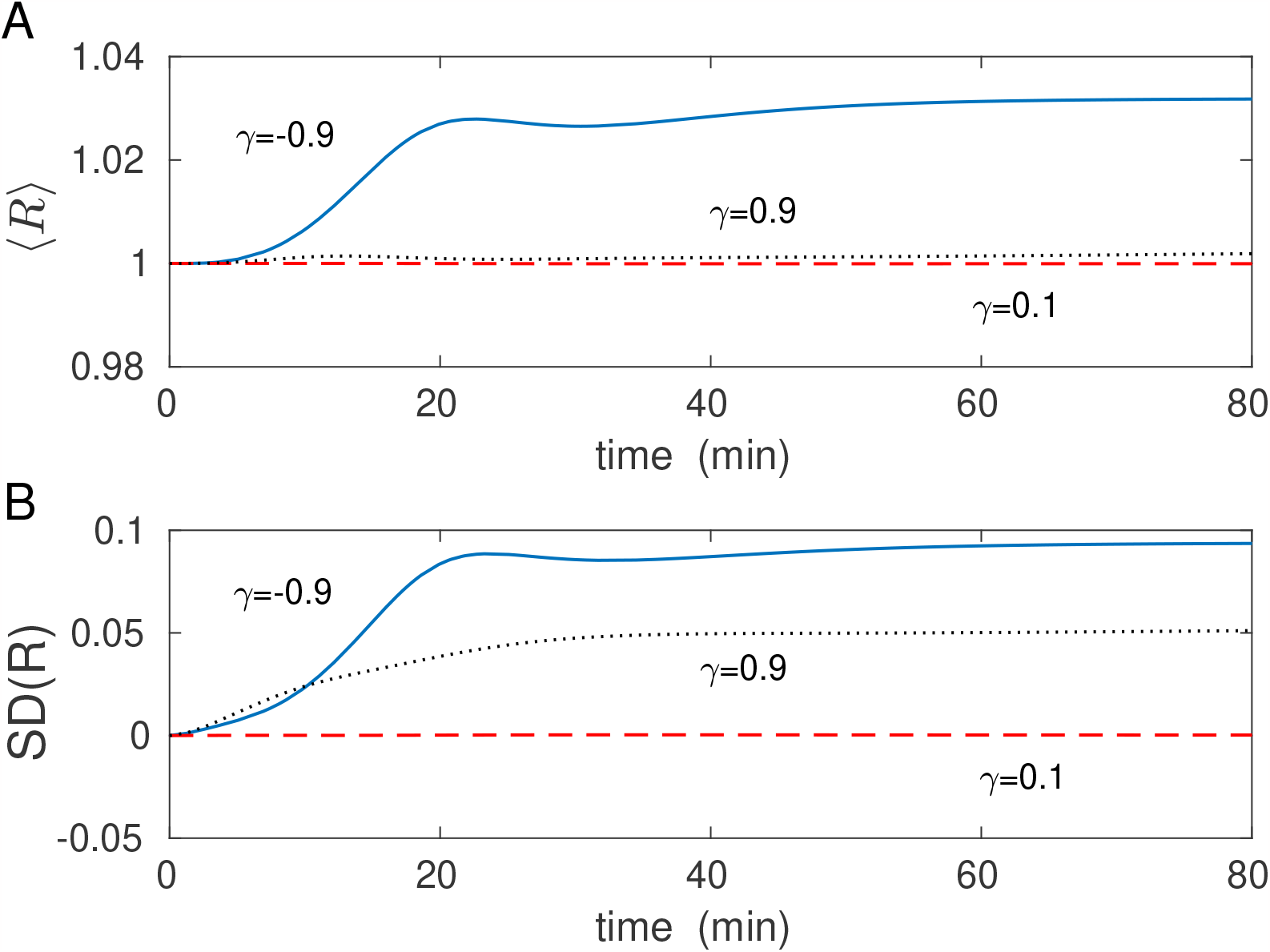
Accuracy measure for pair approximation with N=4 interacting spines. A) Average ratio ⟨*R*⟩ as a function of time for different magnitudes of coupling between the spines. B) Similar as in A but for standard deviation of the ratio *R*. Note that SD(R) is for moderate values of *γ* much less than 0.05, and maximally achieves the value ∼ 0.1 in the extreme case *γ* ↦ −1.

Taken together, the numerical results in Figs. 2 and 3 indicate that the pair approximation derived in this study is quite accurate, and its accuracy is preserved in time. For an analytical example related to the pair approximation, see Appendix B.

## 3. Derivation of entropy production rate as an energy cost for interacting spines

Energy expenditure of synaptic plasticity is associated with transitions between different states of a dendritic spine. The faster the transitions, the more energy is used, and vice versa. Generally, a spine is in a thermodynamic nonequilibrium with its environment, and thus the energy cost is strictly related to entropy production rate of the spine (for general ideas of nonequilibrium thermodynamics, see: Nicolis and Prigogine, (1977), and Peliti and Pigolotti (2021)). Specifically, we assume that the energy rate associated with plasticity processes in dendritic spines is equal to the entropy production associated with stochastic transitions between spine mesoscopic states, similar as in (Karbowski 2019). In our case of *N* dendritic spines, the entropy production rate of the whole system EPR(*s*_1_, …, *s*_*N*_) is given by a general formula for the entropy production (Schnakenberg 1976; Maes et al 2000; Seifert 2012; Van den Broeck and Esposito 2015):

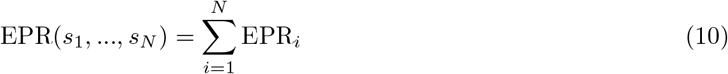

where EPR_*i*_ is the individual entropy productions of each interacting spine

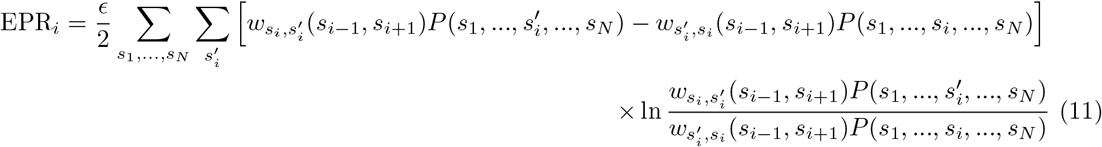

for *i* = 2, …, *N* − 1, and for the boundary terms

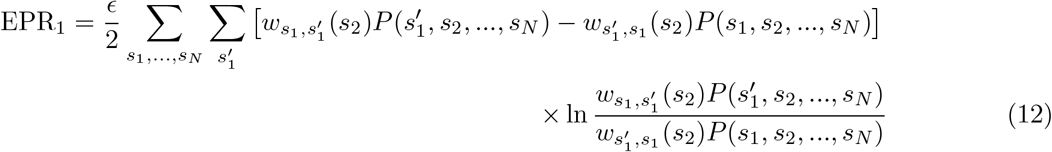

and

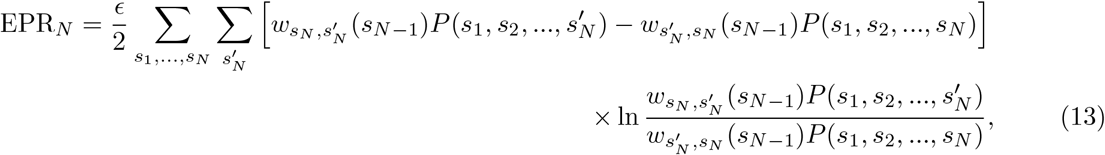

where *ϵ* is the energy scale for various biophysical processes taking place inside a typical dendritic spine and related to molecular plasticity. Its value was estimated at about *ϵ* ≈ 4.6 ·10^5^ kT (or 2.3 · 10^4^ ATP molecules), where *k* is the Boltzmann constant and *T* is the absolute brain temperature (see, Karbowski 2021). In a nutshell, these numbers can be understood by considering that a typical dendritic spine contains roughly ∼ 10^4^ proteins, each with a few degree of freedom corresponding to the number of phosphorylation sites (Sheng and Hoogenraad, 2007).

As before, we explore the pair approximation in Eqs. (11-13), which in the case of *N* spines takes the form

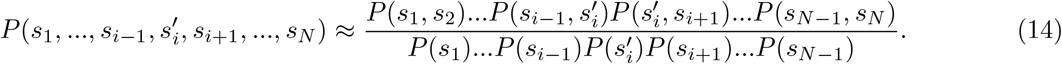

This is a straightforward generalization of formula (6), which can be easily verified. Application of Eq. (14) leads to simplification of the ratio of probabilities under the logarithm in Eq. (11) as

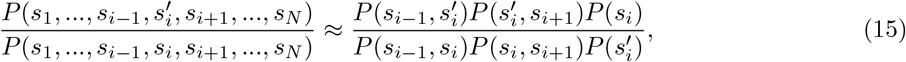

which allows us to perform summation over almost all states *s*_1_, *s*_2_, …, *s*_*N*_ except the few in Eqs. (11-13). This step produces the final expression for the approximate total entropy production rate EPR(*s*_1_, …, *s*_*N*_) of our interacting spines:

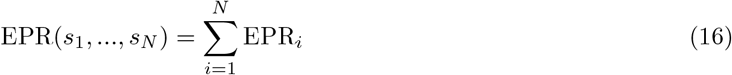

Where

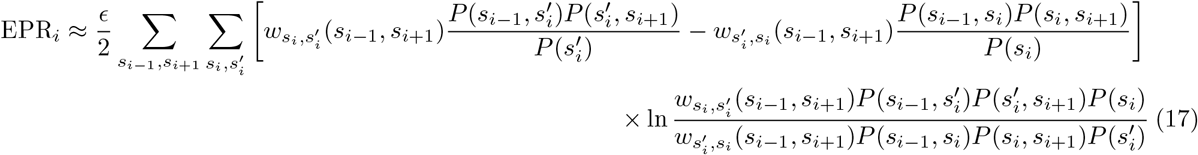

for *i* = 2, …, *N* − 1, and for the boundary terms

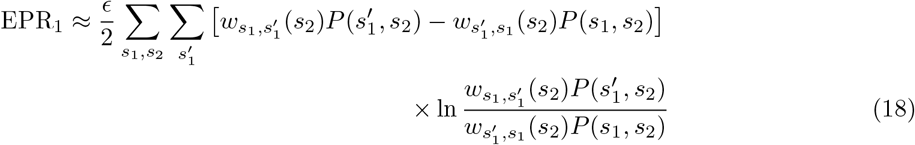

and

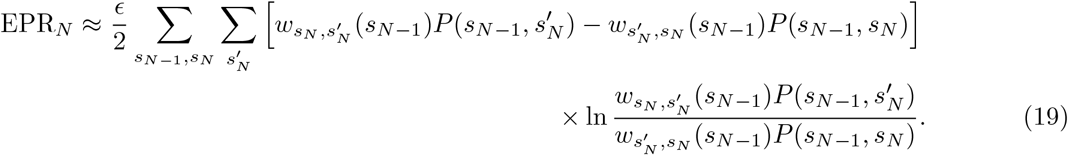

Note that the total entropy production of all spines EPR(*s*_1_, …, *s*_*N*_) is determined exclusively in terms of the two types of probabilities considered in Eqs. (7) and (8), i.e., one- and two-point probabilities. It is also interesting to mention that the form of the approximated EPR in Eq. (17) can be also deduced instantly from the form of the approximated dynamics for probabilities in Eq. (7). This is possible if one realizes that the expression in the bracket on the right in Eq. (7) represents a probability flux.

The total energy *E* used by all spines for LTP induction and its maintenance up to recovery (during synaptic stimulation and post stimulation) is the energy needed to keep the memory trace above the threshold. *E* is the total energy cost of LTP, and it is defined as

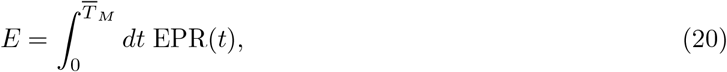

where *t* = 0 relates to the onset of stimulation, EPR(*t*) is the total entropy production rate given by Eqs. (16-19), and 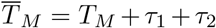, where *T*_*M*_ is the memory time, and *τ*_1_ + *τ*_2_ is the duration of the stimulation (learning phase; see Eq. (3)). The energy used solely for LTP induction and maintenance is denoted as *E*_*ltp*_, and it is given by 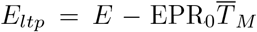, where EPR_0_ is the baseline entropy production rate of all spines.

## 4. Definition of memory trace and memory time

We define the signal associated with dendritic spine activation as

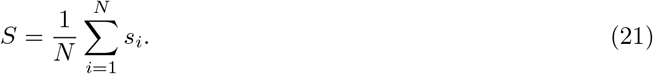

The signal at a steady state (baseline) is denoted as *S*_*ss*_. Memory trace *MT* is defined as the average normalized signal to noise ratio. The normalized signal is simply its deviation from the steady state or baseline. Consequently, the memory trace takes the form:

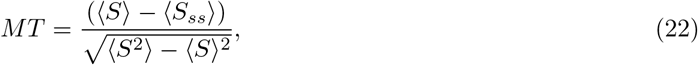

where 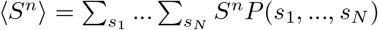 for *n* = 1, 2. The explicit forms for the signals and variance of the signal are

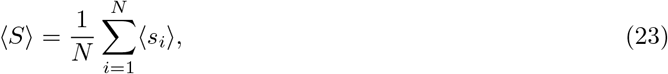

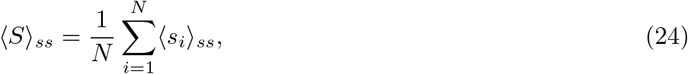

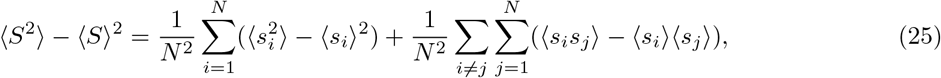

where 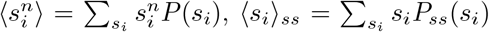, and 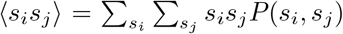, where *P*_*ss*_(*s*_*i*_) is the probability distribution at baseline state. The first sum on the right in Eq. (25) is the sum of variances of individual spines, while the second sum is the total cross correlation of spines. In the pair approximation for the probability, the last sum associated with the correlations simplifies, as only the cross correlations between neighboring spines provide nonzero contributions, since generally *P* (*s*_*i*_, *s*_*j*_) ≈ *P* (*s*_*i*_)*P* (*s*_*j*_) for |*i* − *j*| ≥ 2.

Memory time *T*_*M*_ is defined as the time *t* after stimulation for which memory trace *MT* is in a declining phase and assumes value 1 (Fusi et al 2005; Leibold and Kempter 2008; Karbowski 2019). This is the moment in time when normalized signal becomes comparable to its noise component.

## 5. Derivation of information gain for interacting spines

Information gain *I*, for all spines, right after the end of LTP (i.e., when memory trace decays to the noise level *MT* = 1) is defined as Kullback-Leibler divergence at time 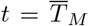, i.e. 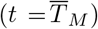, between the baseline steady-state initial probability *P* (*s*_1_, …, *s*_*N*_)_*ss*_ at time *t* = 0 (before LTP stimulation) and final probability at time 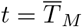, i.e., 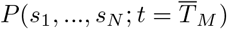. Its form is given by

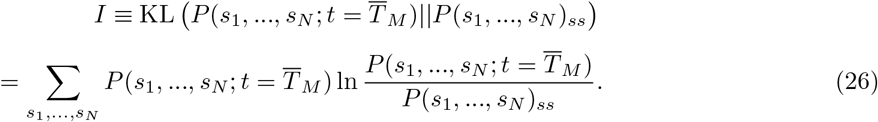

For the pair approximation, i.e., applying the approximation (14) for the probabilities, information gain *I* takes the form

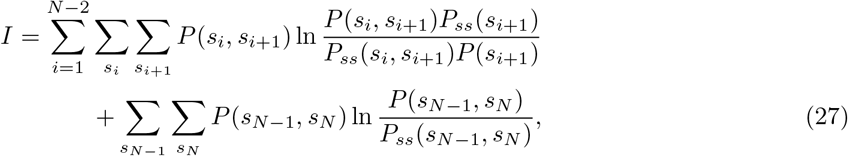

which is used in the computations.

The rate of information gain is equivalent to the rate of Kullback-Leibler KL divergence at arbitrary time *t*, i.e., KL(*t*) (given by Eq. (26) but for arbitrary *t*). We are interested in the rate of information gain, since we want to compare it directly to the entropy production rate, which has a similar information-theoretic meaning. The rate of KL, which we denote as KLR, is given by KLR = *d*KL(*t*)*/dt*. We find

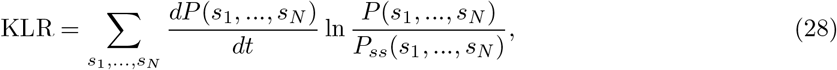

where we used the fact that 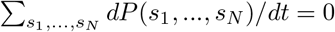. The next step is to substitute Eq. (1) for 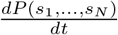, and perform summations over almost all states *s*_1_, …, *s*_*N*_, except the few, similarly to the calculation for EPR. Finally, we use the pair approximation. As a result we obtain the total rate of information gain for all spines as

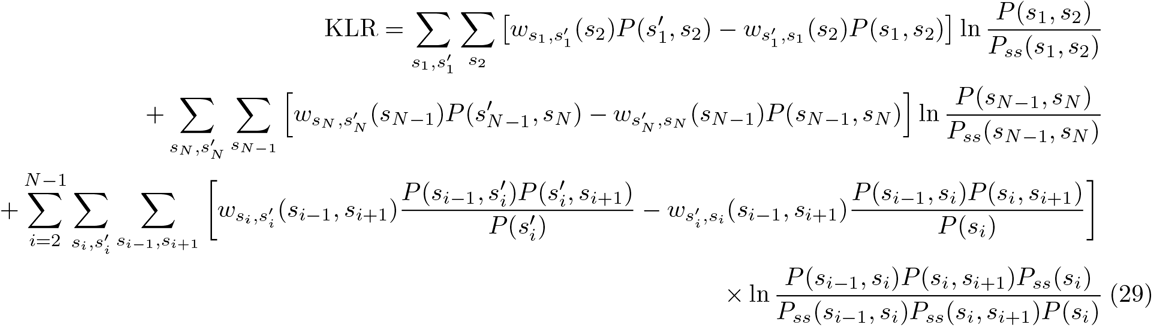

As can be seen, the rate of information gain KLR depends, similar to EPR, on the transition rates between states in all spines. In all figures, we plot KLR per spine, i.e., KLR/*N* . Finally, note that the information gain *I* is the temporal integral of KLR from *t* = 0 to 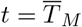, i.e., 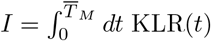.

## 6. Numerical results on memory trace, information gain, and energy cost during LTP

We divide the global dynamics of our system of interacting dendritic spines into two stages. The first stage relates to approaching and reaching a steady-state, starting from random initial conditions (with *f* (*t*) = 0 for each spine). This steady-state is a thermodynamic non-equilibrium steady-state that uses some small but nonzero energy (small but nonzero EPR), and can be thought as the state in which some background information is written in synapses. After reaching the steady-state, we start a second stage associated with spine stimulation, which we call LTP (long-term potentiation) phase. This stage consists of a brief stimulation of the synaptic system by amplifying the transition rates by the function *f* (*t*) present in Eqs. (2) and (3), and then observation of the system recovery to the steady state, with simultaneous recording of the most important observables. Stimulation of synapses is done by turning on the amplifier function *f* (*t*), which amplifies the transition rates between synaptic states (Eqs. 2 and 3). We call this stimulation phase as “learning” phase, and recovery phase as “memory” phase.

### 6.1 Dynamics of memory trace, information and energy rates, and synaptic size associated with LTP induction

Following LTP induction (starting at time *t* = 0 in *f* (*t*) function) memory trace *MT* contained in synapses behaves differently than the rates of information gain (Kullback-Leibler rate KLR) and energy (entropy production rate EPR) (Fig. 4). Initially all three quantities increase sharply, similarly as *f* (*t*), but later their dynamics diverge. Specifically, memory trace exhibits a long temporal tail, i.e., it decays much slower than the stimulation function *f* (*t*), with longer tail for positive cooperation (*γ >* 0) between neighboring spines than for negative cooperation (Fig. 4B). On the other hand, both KLR and EPR (per synapse) decay extremely fast to their baseline values, much faster then *f* (*t*) (Fig. 4C,D). This strongly suggests that keeping memory trace high does not require large rates of energy (EPR). Moreover, the dynamics of KLR and EPR are very similar in shape, and their ratio is positive only initially, when synaptic stimulation increases in time (Fig. 4E). This behavior indicates that information amount written at synapses increases sharply only in the beginning of LTP (learning); at later stages (memory) it weakly decreases (note negative values of KLR/EPR). The rates of information gain (KLR) and energy (EPR) depend weakly, and in the opposite way, on the sign of cooperativity *γ*; the peak of KLR is greater for negative *γ*, while the peak of EPR is greater for positive *γ* (Fig. 4C,D). At its peak, i.e. during stimulation, energy is consumed at rate 4.6 · 10^5^ kT/min, and information is gained at rate 0.1 bits/min, which indicates that acquiring 1 bit at that moment in time is very expensive and costs about 4.6 · 10^6^ kT.

**Fig. 4.**
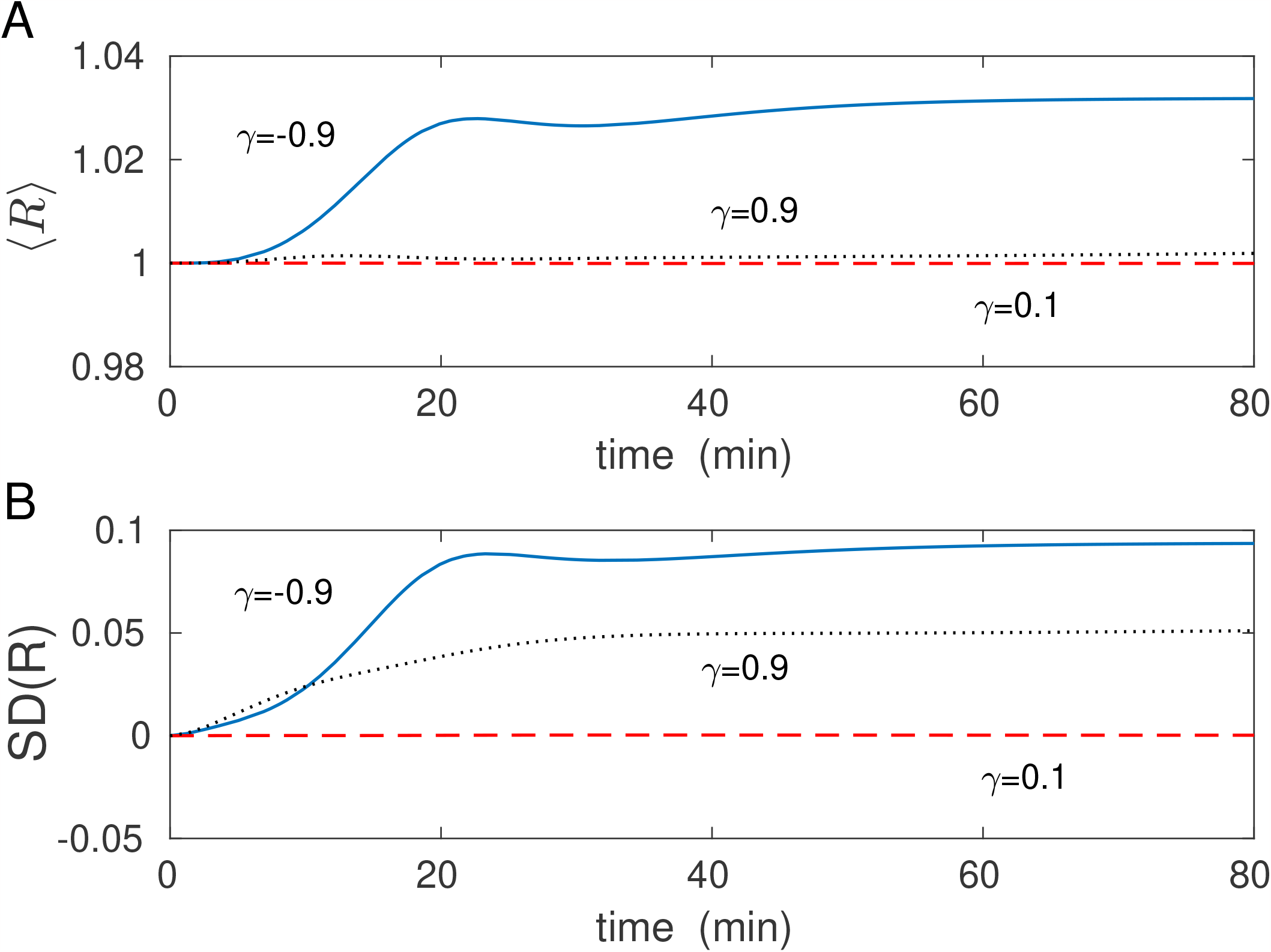
Dynamics of memory trace, and the rates of information gain and energy. Temporal dependence of A) stimulation function *f* (*t*) (amplifier of transitions between the states), B) memory trace MT, C) information gain rate per spine (Kullback-Leibler divergence rate, KLR/*N*), D) entropy production rate (energy rate) per spine (EPR/*N* in units of *ϵ/min*), and of E) the ratio of information gain rate to entropy production rate per spine (KLR/EPR). Learning phase, equivalent to stimulation phase, lasts up to 20 min (A). Memory phase, quantified by memory trace, starts after the end of stimulation and lasts up to ∼ 120 min (B). Note that upon stimulation the rates of information gain and entropy production (KLR and EPR) achieve peaks very fast, but they also decay very fast. In contrast, memory trace lasts much longer. EPR value at its peak is ∼ *ϵ*/min ≈ 4.6 · 10^5^ kT/min per spine, which is about 2-3 orders of magnitude larger than at baseline (before or long after the stimulation).

The dynamics of spine sizes following LTP induction are similar to the behavior of memory trace, except that sizes stabilize at some finite level (Fig. 5). Positive cooperativity among synapses (*γ >* 0) generally indicates positive correlations between them, and vice versa (Fig. 5C). Consequently, positive correlations lead to higher mean spine sizes than negative correlations (Fig. 5B). An opposite effect is seen for the ratio of KLR and spine size; higher peaks are observed for negative cooperativity between synapses (Fig. 5D). Interestingly, normalized correlations are more variable during learning and memory phases for negative cooperativity than for positive one (Fig. 5C).

**Fig. 5.**
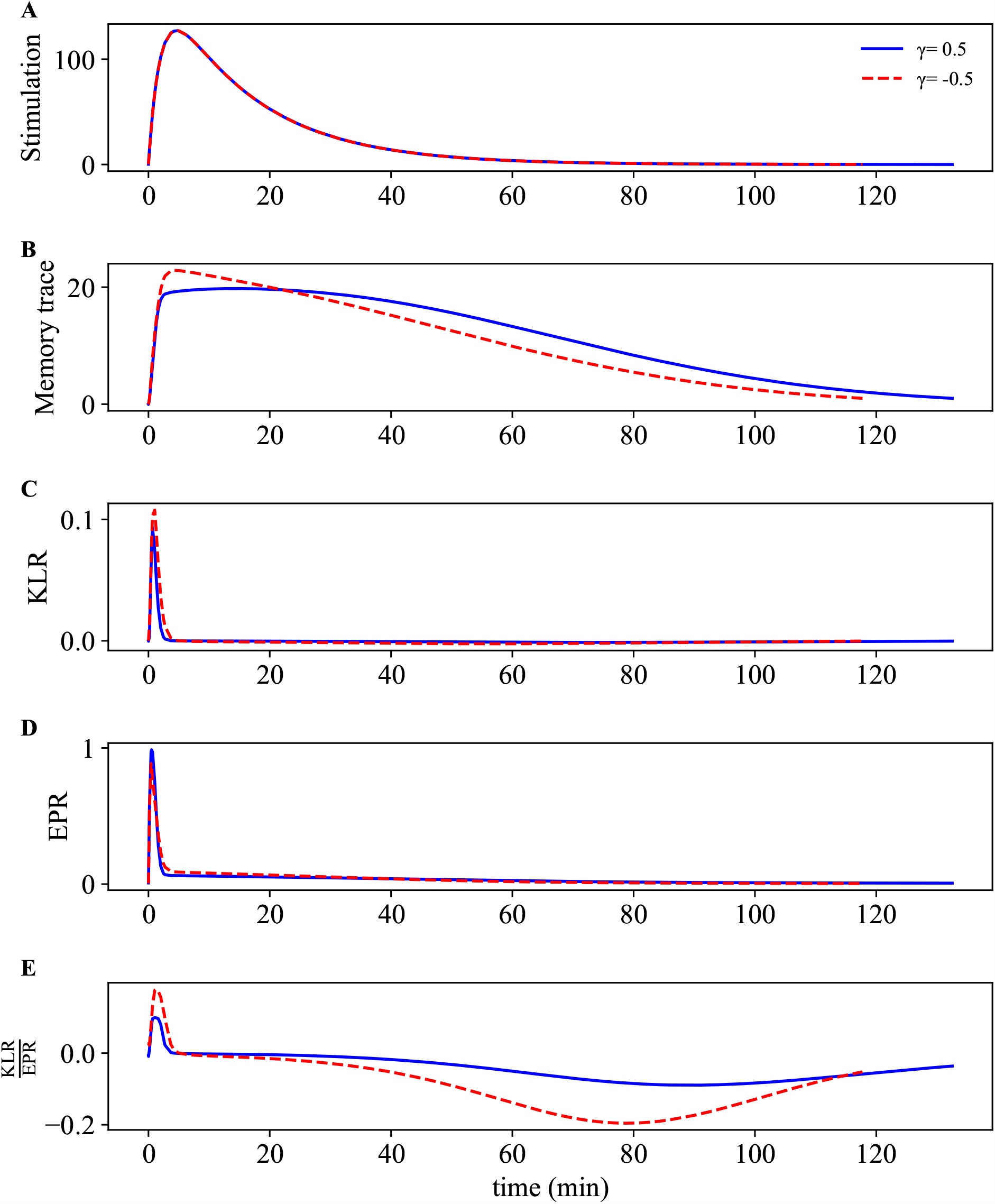
Dynamics of spine size, correlations, and of information gain per spine size. Temporal dependence of A) stimulation function *f* (*t*), B) average spine size ⟨*S*⟩ C) normalized correlations between spines, and of D) the ratio of KLR per spine to average spine size ⟨*S*⟩.

### 6.2 Memory time, spine size, information gain after LTP, and their energy costs as functions of synaptic cooperativity

Memory time *T*_*M*_ grows monotonically but very weakly with cooperativity *γ* up to a point where *γ* is close to its maximal value 1 (Fig. 6A). In that regime of very high positive cooperativity, *T*_*M*_ increases sharply with *γ*. The opposite dependence on *γ* is present for information gain *I* and average spine size (Fig 6B, E). *I* generally decreases with *γ* (for negative *γ* the decay is stronger than for positive *γ*; Fig. 6B), whereas mean spine size monotonically increases with *γ* (increase is stronger for *γ* close to 1; Fig. 6E). Total energy *E* consumed during LTP and its part *E*_*ltp*_ related solely to LTP depend nonmonotonically on cooperativity *γ*, exhibiting broad maxima for *γ* = 0 (Fig. 6C,D). For *γ* close to its maximal value 1, *E* and *E*_*ltp*_ behave in opposite ways: the former increases while the latter decreases with *γ*. This suggests that the cost of LTP alone drops for very high cooperativity.

**Fig. 6.**
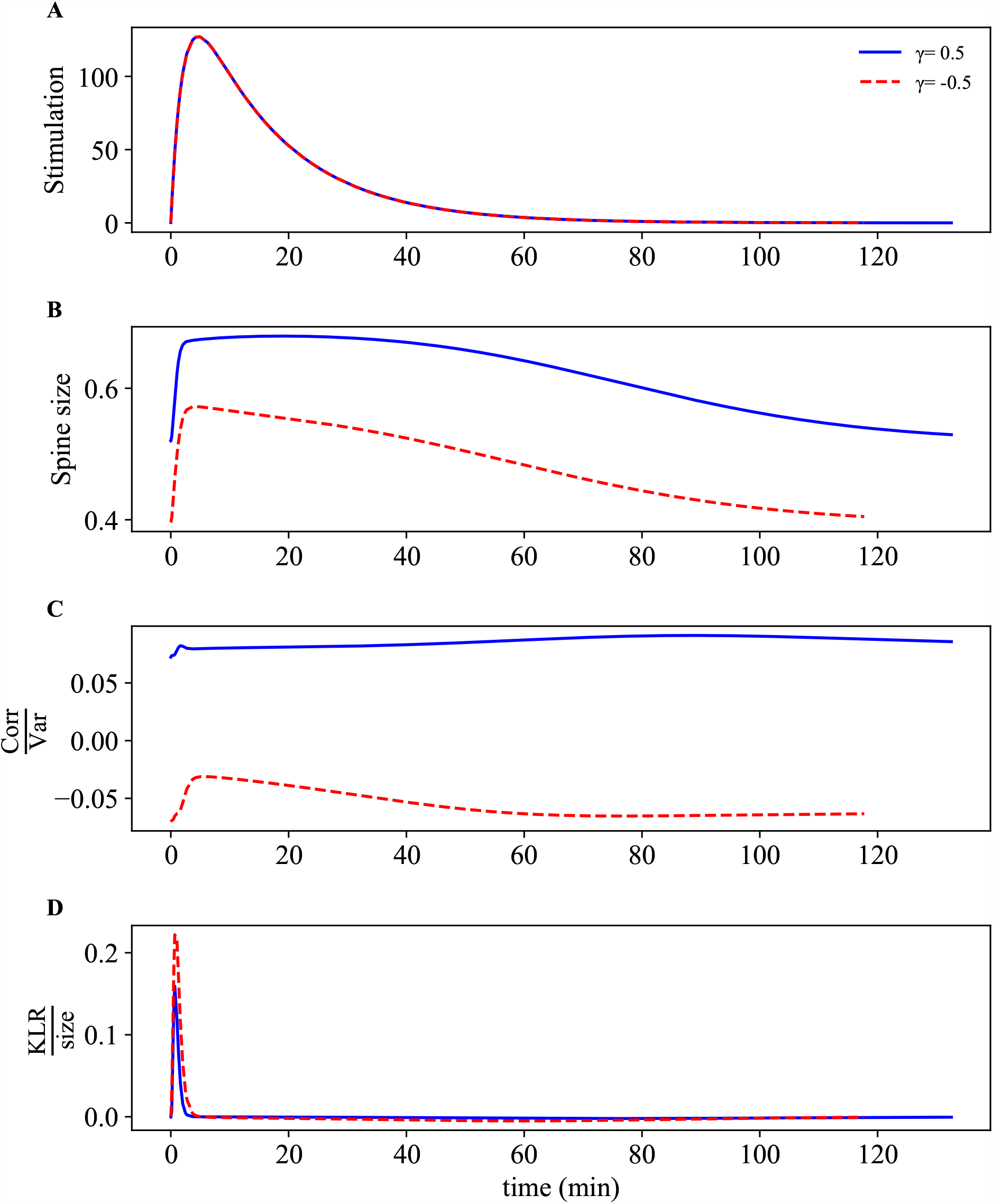
Dependence of memory time, information gain, energy cost, and spine size on the synaptic cooperativity. A) and B) Monotonic but opposite dependence of memory time *T*_*M*_ and total information gain *I* on cooperativity *γ*. C) and D) Total energy consumption of all spines *E* during LTP and its part *E*_*ltp*_, related solely to LTP induction and maintenance, exhibit nonmonotonic dependence on *γ* (energy units are in *ϵ*; see Methods). During whole LTP, a typical spine used about 5*ϵ* ≈ 2.3 · 10^6^ kT of energy. E) Average spine size increases monotonically with *γ*.

How do these results translate to energy and structural efficiency of memory lifetime and information gain? Figure 7 provides the answers. The ratios of memory time and the energies, i.e., *T*_*M*_ */E* and *T*_*M*_ */E*_*ltp*_, are essentially constant for *γ* up to ∼ 0.8 (Fig. 7A), suggesting that memory time and both of these energies grow proportionally with cooperativity for a wide range of *γ*. For larger *γ* these ratios grow significantly with *γ*, especially *T*_*M*_ */E*_*ltp*_, indicating that energy efficiency of memory time is enhanced in the regime of very high positive synaptic cooperativity (Fig. 7A).

**Fig. 7.**
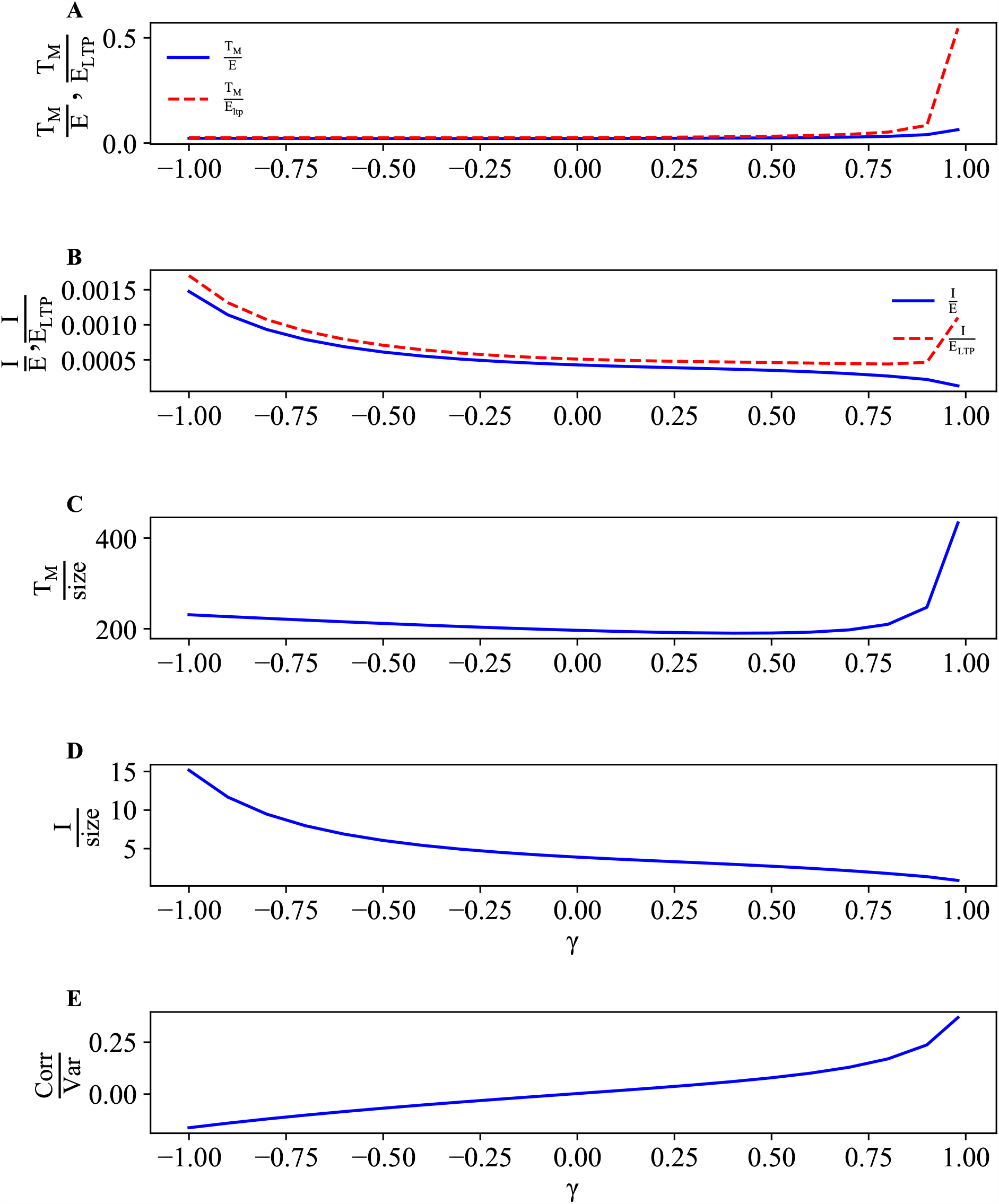
Energetic and structural efficiencies of memory time and information gain in comparison to spine correlations as functions of synaptic cooperativity. Ratios of A) memory time *T*_*M*_ and B) information gain *I* to two energies *E* and *E*_*ltp*_ exhibit two opposite dependence. Similar behavior for the ratios C) memory time and D) information gain to average spine size ⟨*S*⟩ as functions of *γ*. E) Normalized correlations always increase with *γ*, reaching ∼ 0.3 for *γ* ↦ 1.

Energy efficiency of information gain is more complex (Fig. 7B). Generally, the ratio of information to total energy during LTP *I/E* decreases monotonically with *γ*, meaning that efficiency of *I* is maximal for negative cooperativity between synapses. On the other hand, the ratio of information to energy solely to LTP, i.e., *I/E*_*ltp*_ as a function of *γ* has a U-shape, with large values for both high negative and high positive cooperativity. The latter means that information efficiency in that energy currency has two regimes of higher values (Fig. 7B). However, it should be emphasized that the overall energy efficiency of information gain is rather low, at (5 − 10) · 10^−4^ bits/*ϵ* or (1 − 2)·10^−9^ bits/kT, i.e., 1 bit of stored memory (after decline of LTP, i.e. after time ∼ *T*_*M*_) costs about 10^9^ kT (see also below).

Structural efficiency (or energy efficiency of transmission) of memory time and information gain exhibit different behavior as functions of synaptic cooperativity (Fig. 7 C,D). The ratio of information gain to mean spine size *I/*⟨*S*⟩ decreases monotonically with *γ*, which indicates that structural efficiency of information is the largest for negative cooperativity (Fig. 7D), similar to the (total) energy efficiency of *I* (Fig. 7B). The ratio of memory time to mean spine size *T*_*M*_ */*⟨*S*⟩ decreases slightly with increasing cooperativity from negative values of *γ*, but for *γ* ≈ 0.4 this ratio increases sharply with *γ* (Fig. 7C). This result shows that structural efficiency of memory time is the highest for strong positive synaptic cooperativity (Fig. 7C), where spine normalized correlations reach values 0.2 − 0.3 (Fig. 7E).

### 6.3 Sparse representations of synaptic memory and information are more energy efficient

Next we investigate how memory time, information gain, average spine size, and energy cost depend on the fraction *p*_*act*_ of stimulated synapses by presynaptic neurons (Fig. 8). Memory time *T*_*M*_, energy cost solely due to LTP *E*_*ltp*_, and mean spine size grow monotonically with *p*_*act*_, though the first one saturates for larger *p*_*act*_ (Fig 8 A,C,D). In contrast, information gain *I* and total energy cost *E* display non-monotonic behavior: the former has a maximum, while the latter a minimum for a small fraction of active synapses (Fig. 8 B,C). In terms of energy efficiency, the ratios of *T*_*M*_ */E, T*_*M*_ */E*_*ltp*_ and *I/E, I/E*_*ltp*_ have maxima at around the same small fraction *p*_*act*_, regardless of the sign of synaptic cooperativity *γ* (Fig. 9A,B). This means that there exist an optimal percentage of activated synapses on a dendrite that yields the highest information gain and memory lifetime per invested energy (whether total or only due to LTP). For that optimal *p*_*act*_, the energy cost of 1 bit of stored information is about 10^7^ kT, which is much lower (and thus more efficient) than for values *p*_*act*_ away from the optimality. Interestingly, the normalized correlations between spines are essentially independent of *p*_*act*_ (Fig. 9E).

**Fig. 8.**
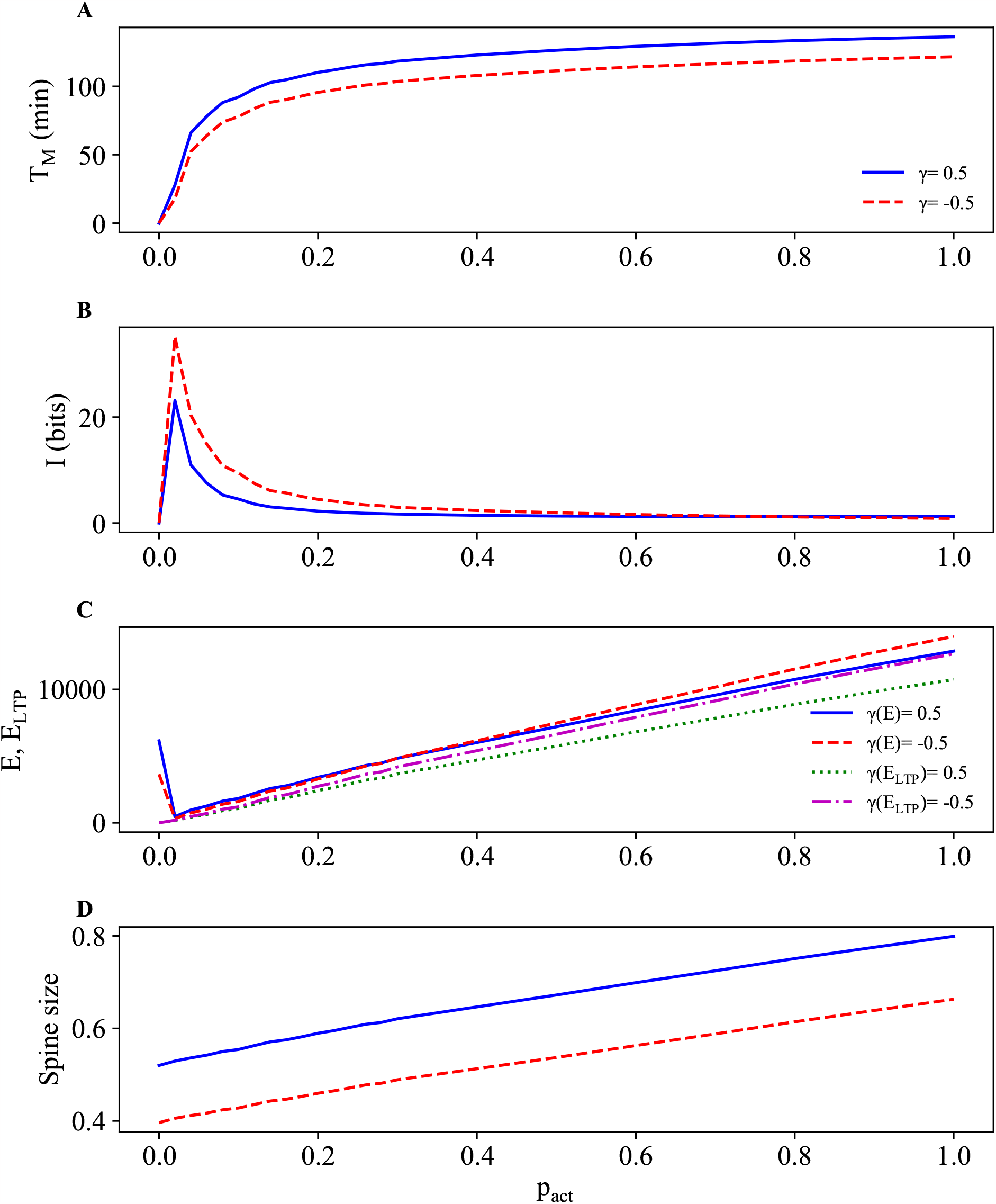
Dependence of memory time, information gain, energy cost, and spine size on probability of synaptic stimulation. A) Memory lifetime *T*_*M*_ saturates for large *p*_*act*_. B) Information gain exhibits a sharp peak for very small *p*_*act*_. C, D) Energy solely due to LTP and average spine size increase linearly with *p*_*act*_.

**Fig. 9.**
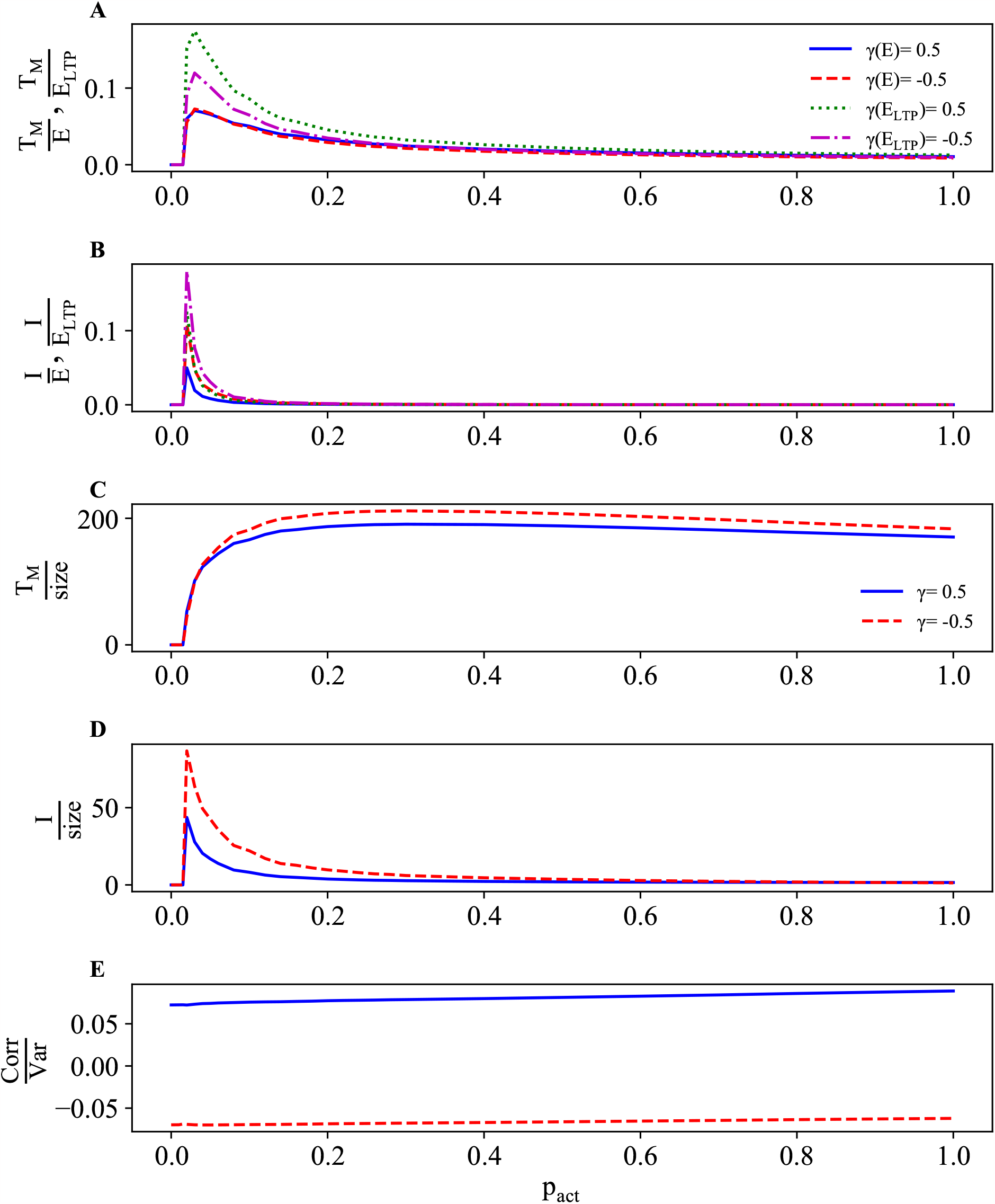
Energetic and structural efficiencies of memory time and information gain as functions of probability of synaptic stimulation. All the ratios of memory time and information gain to energies and to spine size ⟨*S*⟩ exhibit maxima. At the peak, the ratio *I/E* is ∼ 0.05 bit/*ϵ* and *I/E*_*ltp*_ is ∼ 0.15 bit/*ϵ*, which means that 1 bit of stored information after LTP degradation requires about 10^7^ kT of energy.

Structural efficiency of information gain and memory time is qualitatively similar to their energy efficiency (Fig. 9). The information per spine size *I/*⟨*S*⟩ has a similar sharp peak as *I/E* for low *p*_*act*_ (Fig. 9D). However, memory time per spine size *T*_*M*_ */*⟨*S*⟩. has a much broader maximum at higher values of *p*_*act*_ (Fig. 9C).

Taken together, these results indicate that energetic and structural efficiency of information and its duration in synapses can be optimized for low fractions of activated synapses during LTP. In other words, acquiring and storing of synaptic information can be most efficient by using sparse synaptic representations, regardless of the nature of synaptic cooperativity.

### 6.4 Memory time, spine size, information gain, and their energy costs as functions of strength and duration of stimulation

In Figs. 10 and 11, we show how memory time, information gain, energy cost, and mean spine size depend on the duration of stimulation *τ*_1_ (decay time of the stimulation). Memory time *T*_*M*_ and its energy costs *E, E*_*ltp*_ both grow proportionally with *τ*_1_ (Fig. 10A,C), such that their ratios *T*_*M*_ */E* and *T*_*M*_ */E*_*ltp*_ are almost constant, although with a weak increasing trend (Fig. 11A). Information gain *I* as well as its energy efficiencies *I/E, I/E*_*ltp*_ decrease with *τ*_1_ for small *τ*_1_, but for larger *τ*_1_ all of these quantities saturate at some small level (Figs. 10B and 11B). Average spine size shows a similar saturation effect after a small initial increase (Fig. 10D). In terms of structural efficiency, *T*_*M*_ */*⟨*S*⟩ grows linearly with *τ*_1_ (Fig. 11C), and *I/*⟨*S*⟩ first decreases with *τ*_1_ and then saturates (Fig. 11D). The results in Fig. 11 indicate that longer stimulation times are not particularly beneficial for energy efficiency of memory lifetime and information gain (Fig. 11A,B). However, longer stimulation may be advantageous for structural efficiency of memory duration (Fig. 11C), though not for information *I* (Fig. 11D).

**Fig. 10.**
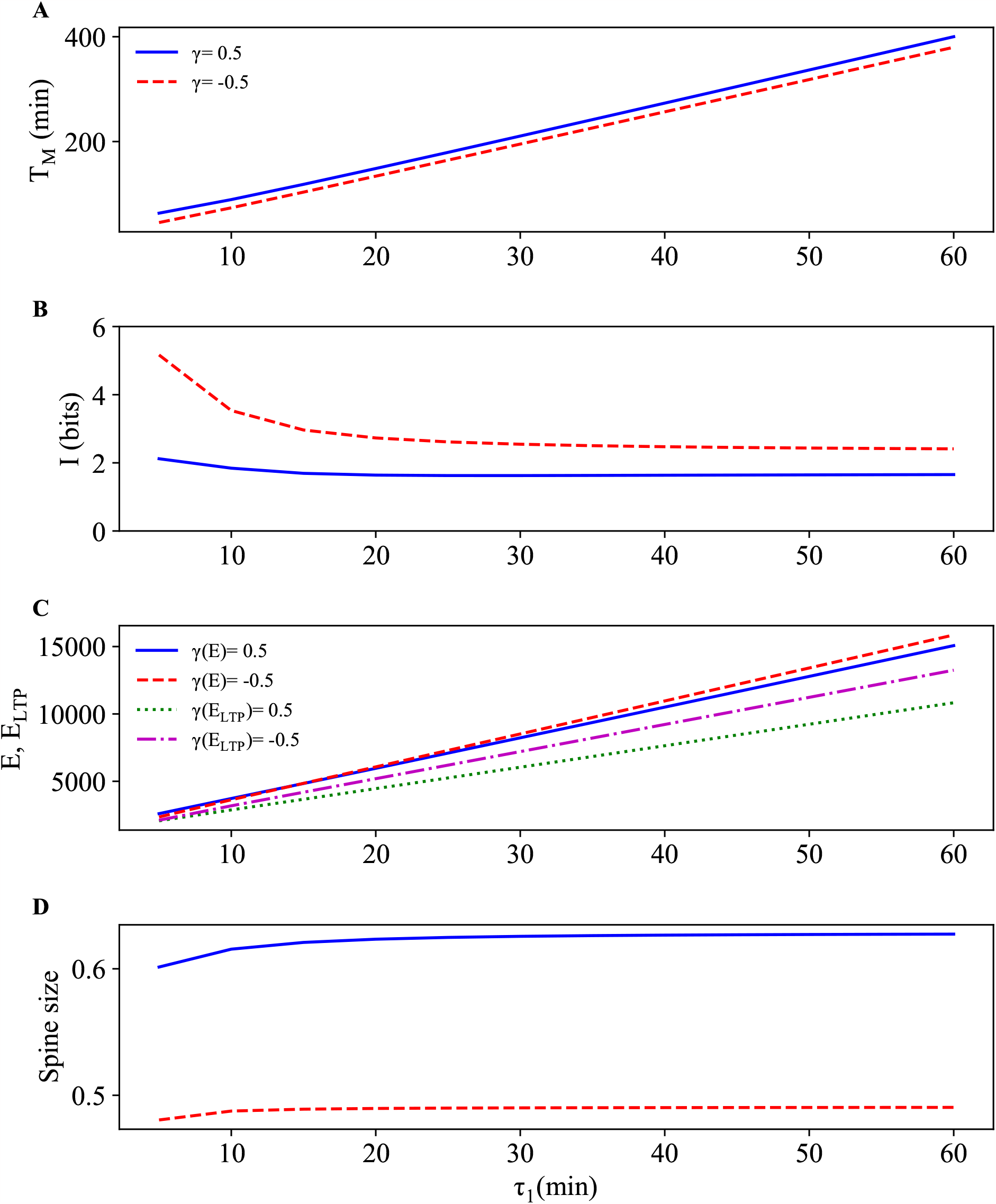
Dependence of memory time, information gain, energy cost, and spine size on duration of stimulation *τ*_1_.

**Fig. 11.**
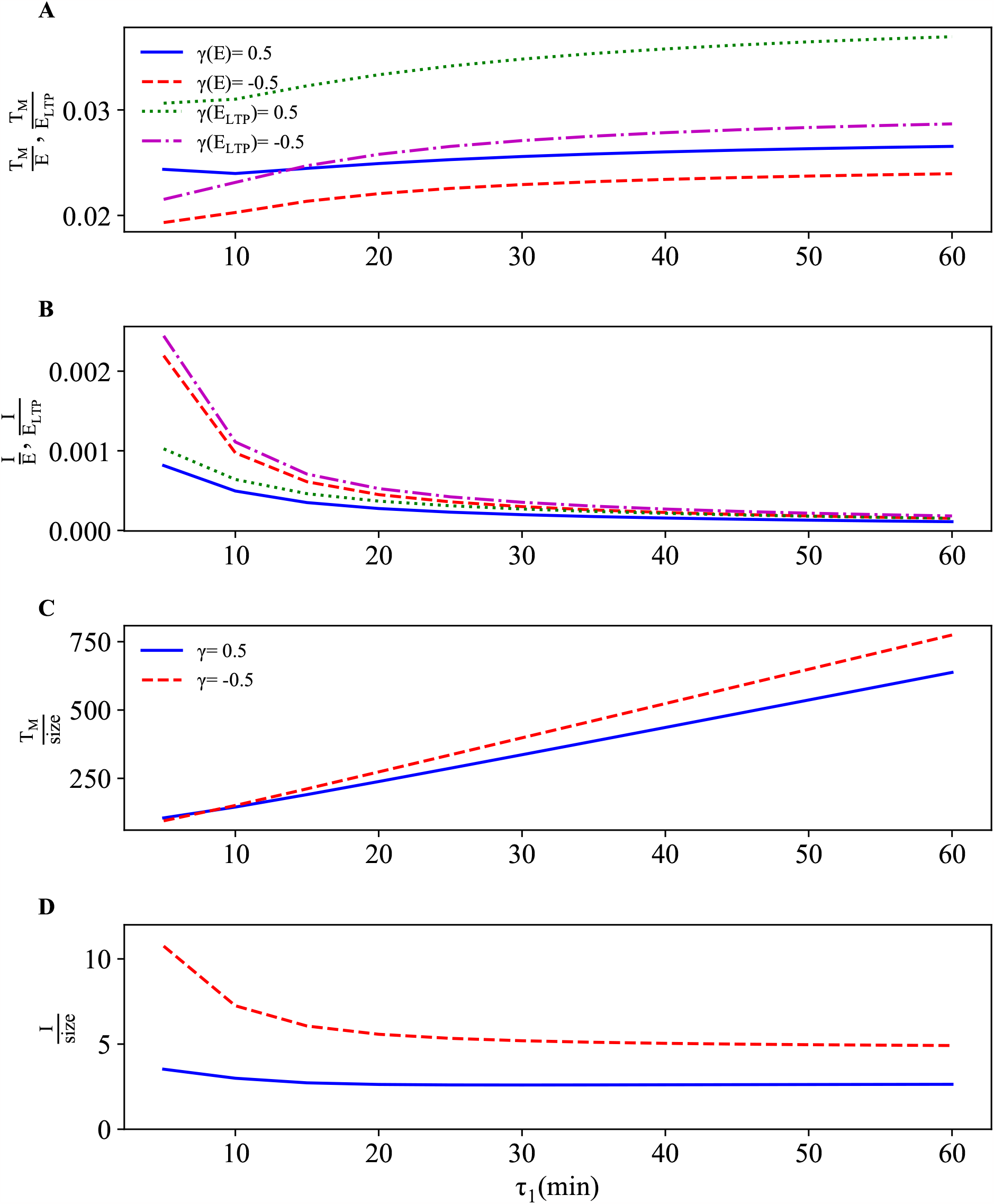
Energetic and structural efficiencies of memory time and information gain as functions of duration of stimulation *τ*_1_. Note that energetic efficiency drops with increasing the duration of stimulation (B). Structural efficiency of memory lifetime grows linearly with *τ*_1_ (C).

In Figs. 12 and 13, we present the dependence of memory time, information gain, energy cost, and mean spine size on the amplitude of stimulation *A*. Memory time *T*_*M*_ and energy costs both grow weakly but saturate with *A* (Fig. 12 A,C). On the other hand, *I* and spine size ⟨*S*⟩ stay almost constant (Fig. 12 B,D). These results translate into very weak variability of energy and structural efficiencies of memory time and information gain on *A*, which are close to constancy, except the ratio *T*_*M*_ */*⟨*S*⟩ that exhibits an increasing trend, but with a saturation (Fig. 13). This suggests that too strong stimulations are also not advantageous over weaker stimulations for efficiency of information gain and its duration.

**Fig. 12.**
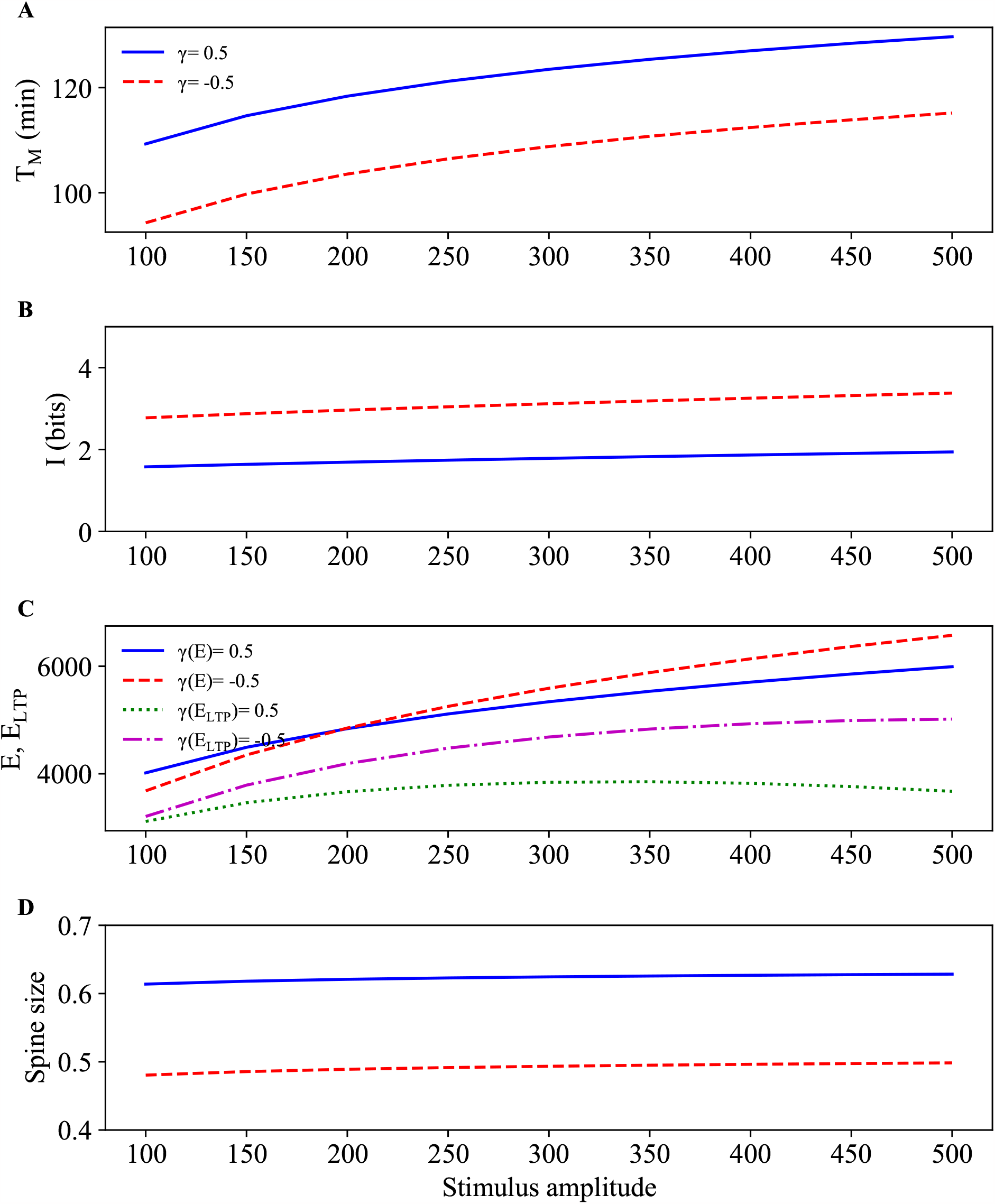
Dependence of memory time, information gain, energy cost, and spine size on strength of stimulation A. Note that information gain (A) and mean spine size (D) are essentially independent of *A*.

**Fig. 13.**
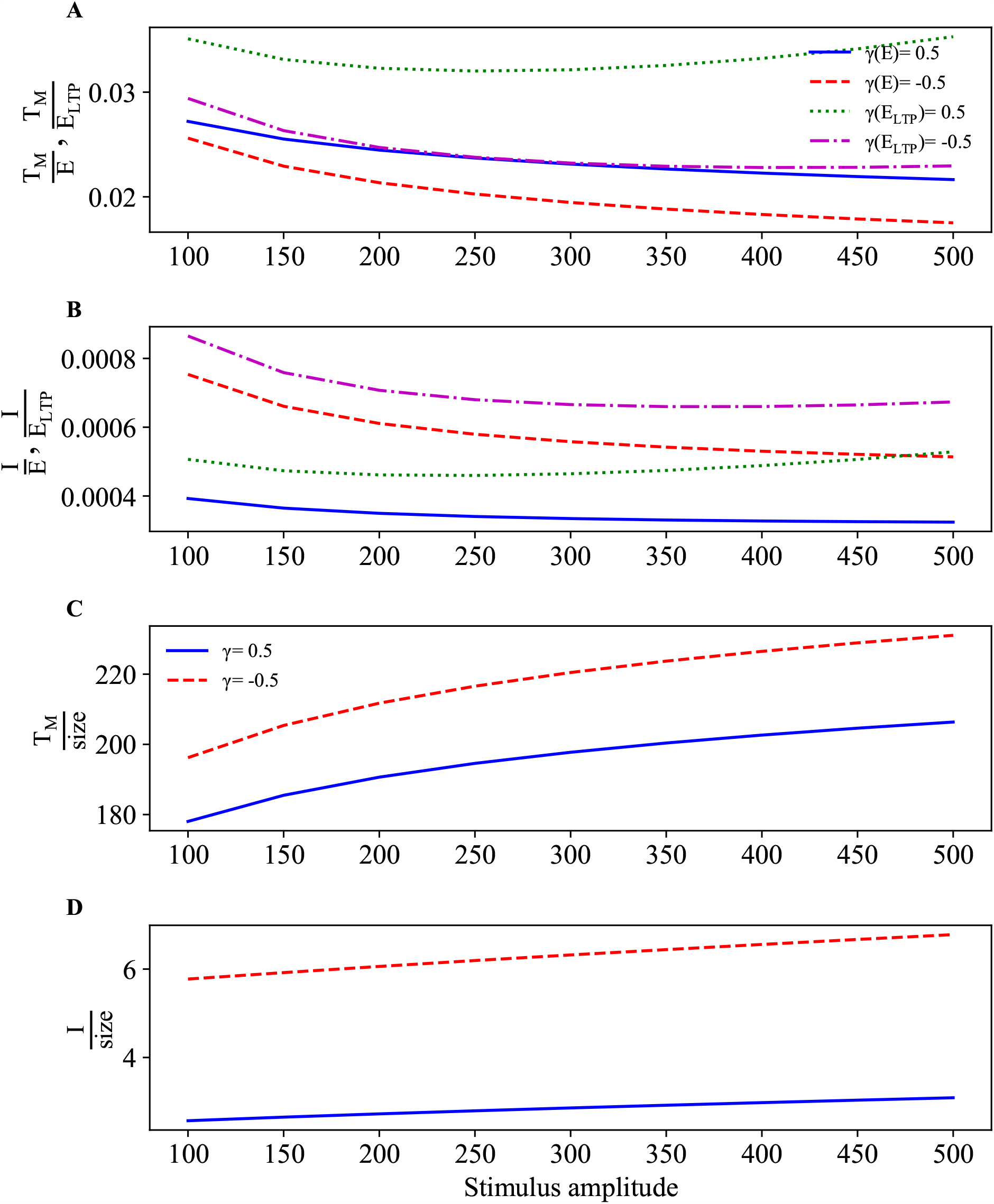
Energetic and structural efficiencies of memory time and information gain as functions of strength of stimulation A. Note that too large stimulation amplitudes are not beneficial for the energy and structural efficiencies (saturation effects).

### 6.5 Influence of synaptic number on the efficiencies of memory lifetime and information gain

In Fig. 14, we show that energy efficiency of both memory lifetime *T*_*M*_ and information gain *I* exhibit a decreasing trend with increasing the number of dendritic spines *N*. The biggest drop in efficiency is for changing *N* from 10 to ∼ 2000 (Fig. 14A,B). For higher values of *N*, the rate of decline is much slower. This result suggests that having large number of spines on a dendrite is generally highly inefficient in terms of energy for storing information. For example, for *N* = 10 we obtain *T*_*M*_ */E* ≈ 1 min/*ϵ*, and *I/E* ≈ 4 ·10^−2^ bits/*ϵ*, i.e., storing 1 min of memory in the spines costs about *ϵ* ≈ 4.6 · 10^5^ kT of energy, and storing 1 bit after the degradation of LTP costs 25*ϵ* ≈ 10^7^ kT. In contrast, for *N* = (8 − 10) · 10^3^, we have *T*_*M*_ */E* ≈ 2 · 10^−3^ min/*ϵ* and *I/E* ≈ 10^−4^ bits/*ϵ*, which means that in this case storing 1 min of memory costs ∼ 500*ϵ* ≈ 2 · 10^8^ kT, and storing 1 bit in all *N* spines after LTP degradation costs 10^4^*ϵ* ≈ 4.6 · 10^9^ kT.

**Fig. 14.**
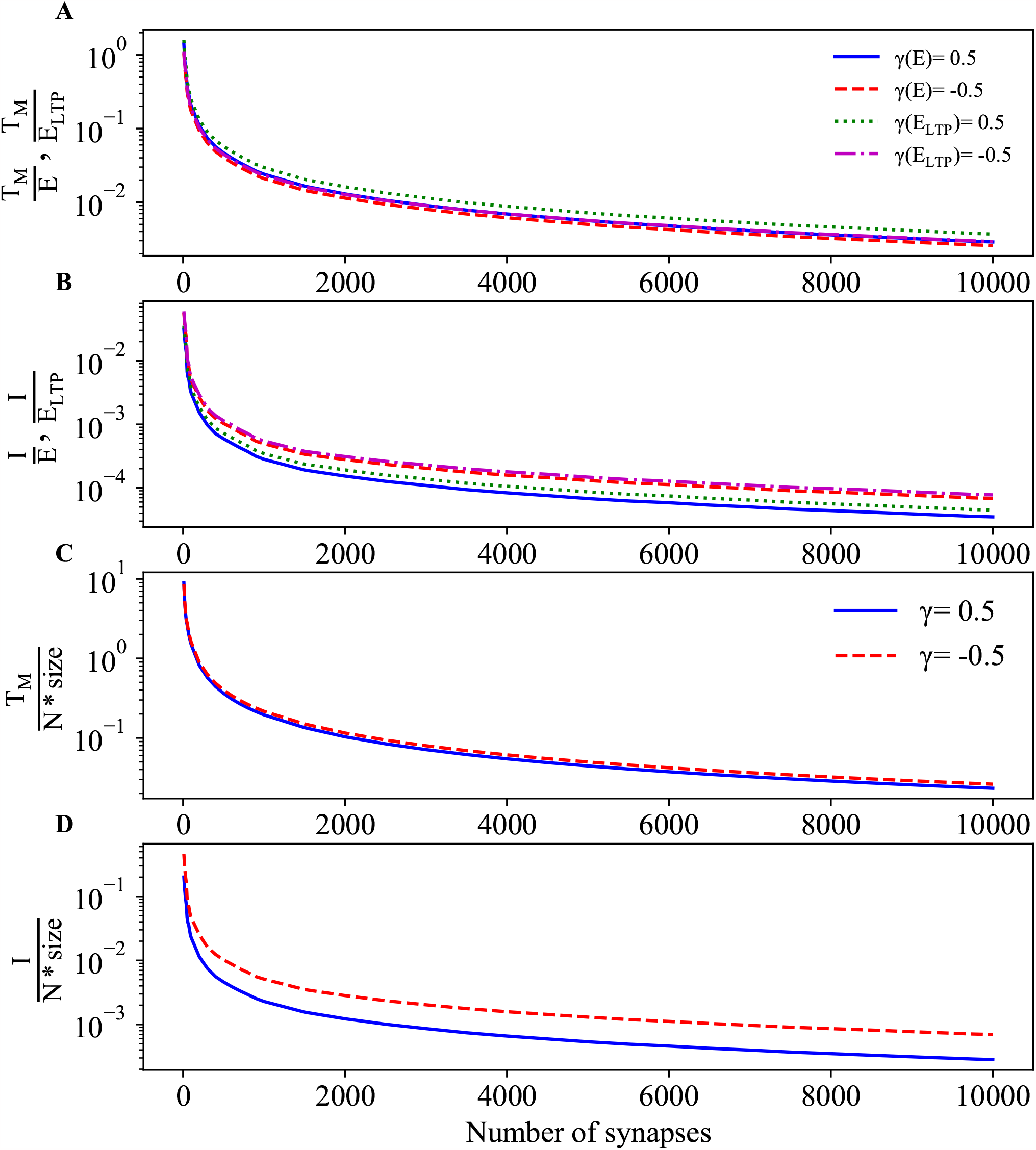
Energy and structural efficiencies of memory lifetime and information gain drop with increasing synaptic numbers *N* . (A, B) Energy efficiency of *T*_*M*_ and *I* decreases dramatically, by two orders of magnitude, with increasing the number of spines from *N* = 10 to *N* = 2000, and much slower with increasing the number of spines from *N* = 2000 to *N* = 10000. (C, D) Essentially the same declining effect is observed for structural efficiency of *T*_*M*_ and *I*, i.e., *T*_*M*_ */*(*N* ⟨*S*⟩) and *I/*(*N* ⟨*S*⟩).

Since memory lifetime and information gain are collective variables, to determine structural efficiencies of these variable as functions of *N*, we have to divide *T*_*M*_ and *I* by the whole structural cost of all spines, i.e., by *N* · ⟨*S*⟩. Thus, the structural efficiencies in this case are the ratios *T*_*M*_ */*(*N* · ⟨*S*⟩) and *I/*(*N* · ⟨*S*⟩) (Fig. 14 C,D), and they drop significantly with increasing *N*, in a similar manner as the corresponding energy efficiencies (in Fig. 14 A,B).

Taken together the two types of the efficiency, it is clear that too many synapses on a dendrite is not beneficial for information storage and its duration in terms energy and biochemical resources.

## 7. Summary and Discussion

We formulated the probabilistic approach to global dynamics of interacting synapses (on a single dendrite) and to their nonequilibrium thermodynamics to study information processing in synaptic internal degrees of freedom. In order to make the high dimensional system of interacting synapses computationally tractable, we introduced the so-called pair approximation, which effectively reduces the dimensionality and number of equations describing the system dynamics in a closed form by considering only the probabilities of singlets and doublets of dendritic spines. We verified on a simplified example that this approximation provides a very good accuracy of the exact dynamics, as well as entropy production rate. We also gave the analytical condition for the applicability of the pair approximation in the equilibrium Ising model in Appendix B (for a more general discussion about the accuracy of the pair approximation, see Matsuda et al (1992), and van Baalen (2000)).

The Master equation approach combined with stochastic thermodynamics allows us to treat information contained in synaptic states on equal footing with its energy cost by relating them to, respectively, the rates of Kullback-Leibler divergence and entropy production. Both of these quantities depend on state probabilities as well as on transitions between the states, and we provided explicit formulas for their calculations. The formalism makes it clear that every plastic transition in synaptic system is associated both with some information processing (flow) and with some energy expenditure (entropy production). Even stationary states out of thermodynamic equilibrium (baseline states), in order to maintain them, require some energy (usually small), which essentially means that keeping information always costs some energy for nonequilibrium systems even in stationary conditions. Physically, this energy dissipation (entropy production) in stationary state is a consequence of breaking the so-called detailed balance in probability flows, and it is a generic feature of systems out of thermodynamic equilibrium (in the context of physics, see: Maes et al 2000; Seifert 2012; for a dendritic spine, see: Karbowski 2019).

Our main results are: (i) Learning phase of a signal (spine stimulations) involves high levels of both information and energy rates, which are much larger than their values during a memory phase (Fig. 4). This indicates that keeping information (memory) is relatively cheap in comparison to acquiring it (learning). This result on a level of many interacting synapses is qualitatively similar to the result in (Karbowski 2019), where it was shown that memory trace resulting from molecular transitions in a single synapse (protein phosphorylation) decouples from energy rate after stimulation phase, leading to a relatively cheap long memory trace of protein configurations. In our model, the maximal energy rate of spine plasticity taking place during stimulation is ∼ 4.6 · 10^5^ kT/min (for *p*_*act*_ = 0.3). (ii) Memory lifetime and its energy efficiency can significantly increase their values for very strong positive synaptic cooperativity, while the opposite is observed for information gain right after LTP drops to its noise level (Fig. 7). This result suggests that strong local positive correlations between neighboring spines can be beneficial for memory storage, especially in the range ∼ 0.3 (Fig. 7E), i.e., for values reported experimentally (Makino and Malinow 2011). This conclusion supports the so-called “synaptic clustering hypothesis”, which was proposed as a mechanism for producing synaptic memory (Govindarajan et al 2006) and enhancing its capacity (Poirazi and Mel 2001; Kastellakis and Poirazi 2019). (iii) There exists an optimal fraction of stimulated synapses during LTP for which energy efficiency of both memory lifetime and information gain exhibit maxima (Fig. 9). This means that sparse representations of learning and memory are much better in terms of energy efficiency, and thus might be preferable by actual synapses. This is also true for structural efficiency of information gain (Fig. 9). (iv) Energy and structural efficiencies of memory lifetime and information gain after degradation of LTP both drop dramatically with increasing the number of spines (Fig. 14). For example, storing 1 bit after LTP is over costs “only” ∼ 10^7^ kT for 10 spines, and at least two orders of magnitude more, i.e., ∼ 4.6 · 10^9^ kT for ∼ 10^4^ spines. Interestingly, the latter figure for energy cost is about ∼ 10^4^ times larger than the cost of transmitting 1 bit through a chemical synapse (Laughlin et al 1998). This high inefficiency of *T*_*M*_ and *I* is surprising and can induce a question: why do neurons have so many synapses? Perhaps, the answer is that neurons need many synapses for reliable transfer of an electric signal (short-term information) between themselves, not necessarily for efficient storing of long-term information.

Our model is based on the assumption that a dendritic spine can be treated as the system with discrete states, which is compatible with some morphological observations (Bourne and Harris 2008; Montgomery and Madison 2004; Bokota et al 2016; Urban et al 2020). In this respect, it is similar in architecture to some previous discrete models of synapses or dendritic spines (Fusi et al 2005; Leibold and Kempter 2008; Barrett et al 2009; Benna and Fusi 2016). However, these models treat synaptic states quite arbitrary and abstract, and consider mostly unidirectional transitions between the states, which makes these models thermodynamically inconsistent (e.g. entropy production rate, equivalent to plasticity energy rate, is ill-defined and yields infinities for unidirectional transitions). In contrast, our model takes as a basis well defined morphological synaptic states, with bidirectional transitions between them that are estimated based on empirical data (Bokota et al 2016; Basu et al 2018; Urban et al 2020). The latter feature, i.e., bidirectional transitions, makes our model thermodynamically consistent (with finite entropy production), as was explained in a previous model of metabolic molecular activity in a single spine within the framework of cascade models of learning and memory (Karbowski 2019).

In this paper, we consider two types of costs. The first is energy cost (related to entropy production) associated with stochastic “plastic” transitions between different synaptic states. The second is structural cost, defined here as proportional to average spine size and related to biochemical cost of building a synapse. This structural cost is also proportional to the rate of electric energy of synap-tic transmission (Attwell and Laughlin 2001; Karbowski 2009, 2012), as spine size is proportional to synaptic electric conductance or, more commonly, synaptic weight (Kasai et al 2003). For standard cortical conditions, i.e. for low firing rates ∼ 1 Hz, the energy cost of synaptic transmission is much larger than the energy cost related to plasticity processes inside spine (Karbowski 2019). However, these two costs can become comparable for very large firing rates, ∼ 100 Hz (Karbowski 2021). There is some confusion in the literature about these two types of energy costs, and some researchers associate the structural cost related to synapse size/weight with “plasticity energy cost”, by assuming the larger synaptic weight leads the higher plastic energy cost (e.g. Li and van Rossum 2020). However, this does not have to be so, and we should make a distinction between the two energy costs. Bigger synapses do not necessarily require larger amounts of plasticity related energy than smaller synapses, because the transitions in bigger synapses could be generally much slower than in smaller synapses (as is in fact reported in some experiments; Kasai et al 2003). Thus, what mostly matter for the plasticity energy rate are the rates of transition between internal synaptic states. On the other hand, synaptic size/weight is always a good indicator of electric energy rate related to fast synaptic transmission (Attwell and Laughlin 2001; Karbowski 2009, 2012).

The present model can be extended in several ways, e.g., by introducing heterogeneity in spine interactions. That is, by allowing random signs of the cooperativity parameter *γ*. However, we suspect that such modifications would not alter the general qualitative conclusions. Finally, our model considers only the early phase of LTP, the so-called e-LTP, which generally lasts up to a few hours and does not involve protein synthesis inside spines. Inclusion of protein synthesis, associated with the process of memory consolidation and thus late phase of LTP (so-called l-LTP), would require some modifications in our present model. The most important of which is inclusion of additional variables in the probabilities characterizing spine states, which are related to internal degrees of freedom (proteins, actin, etc).

This clearly would make the model much more complex, and thus it remains a major challenge at present. However, we hope that our approach of stochastic thermodynamics provides some insight about attempting to model the interplay of information and energy during the late phase of LTP and memory consolidation for interacting spines.

Appendix A: Stochastic model of morphological states in dendritic spines.

Our data on dendritic spines come from cultures of rat hippocampus (Bokota et al 2016; Basu et al 2018). We assume that each dendritic spine can be in four different morphological states: nonexistent (lack of spine), stubby, filopodia, and mushroom. These four states constitute the minimal number of states that can be classified and quantified on a mesoscopic level (Bokota et al 2016; Basu et al 2018; Urban et al 2020). Each state has a typical size, which can be characterized by several geometric parameters. We focus on one particular parameter, spine head surface area, as an indicator of both spine structure and function. Spine head surface area is proportional to synaptic weight (as measured by the number of AMPA receptors; Kasai et al 2003), which relates to spine neurophysiological function (synaptic transmission and information storage in molecular structure). On the other hand, spine area is also a measure of its structural and metabolic costs (larger area larger both costs). To estimate spine areas in each of the three states (for nonexistent state the size is 0), we used the data on minimal and maximal spine head diameters from (Bokota et al 2016; Urban et al 2020), which gave us the following numbers: *d*(0) = 0 for nonexistent, *d*(1) = 0.496 *μ*m^2^ for stubby, *d*(2) = 0.786 *μ*m^2^ for filopodia/thin, and *d*(3) = 1.045 *μ*m^2^ for mushroom. Values of these intrinsic transition matrix are given in Table 1. The values of global parameters are presented in Table 2.

We assume that global spine dynamics can be described as Markov chain model, in which there are stochastic transitions between spine internal states. The general model of this kind is given by the following master equation (e.g. Glauber 1963):

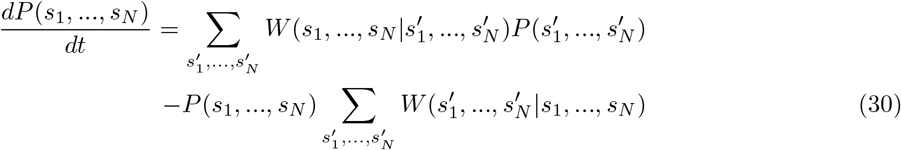

where 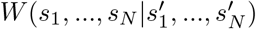 is the multidimensional transition matrix of the whole system of *N* dendritic spines. We assume that transitions between the states take place only in one of the spines at any given time unit (the rest of states in other spines do not change in that brief time step). This means that the multidimensional matrix 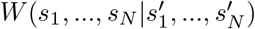 can be decomposed as

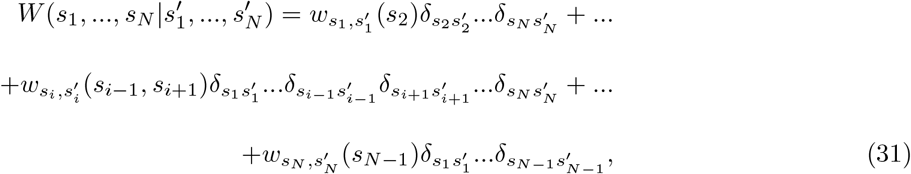

where 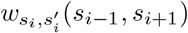 are the transitions rates at individual spines, which are dependent on the neighboring spines. After insertion of the form of multidimensional transition matrix *W* in Eq. (31) above, and performing summations, we obtain Eq. (1) in the main text.

By using the above pattern of transition probabilities we assume that at sufficiently short time step only one transition can take place, i.e., simultaneous transitions in different spines are much less likely, and thus can be neglected. Indeed, since the local basic transitions between mesoscopic states in individual spines are of the order of several minutes (Urban et al 2020; Table 1), the likelihood that two or more such transitions in two or more spine take place simultaneously in a short period of time (much smaller than a minute) is small. This type of locality of explicit synaptic interactions allows us to analyze the dynamics of global system of *N* interacting spines.

Appendix B: Validity of the “pair approximation” for analytically solvable model

In this section we provide conditions that much be satisfied for applying the pair approximation in a case that can be treated analytically, which is a simplified Ising model in thermal equilibrium. We consider 3 interacting units (*i* = 1, 2, 3) forming a linear ordered chain, similar as in Fig. 1, but each unit having only two states *s*_*i*_ = −1 or *s*_*i*_ = 1. Additionally, in this model, nearest neighbors interact strongly with the coupling *J*, whereas remote units (i.e. 1 and 3) interact weakly with the coupling *κ*, which is much smaller than *J*. Our goal is to check how accurate is the pair approximation as we increase the strength of remote coupling *κ* in relation to *J*.

The equilibrium probability of finding a given configuration of units *s*_1_, *s*_2_, *s*_3_ has the form (Feynman 1972)

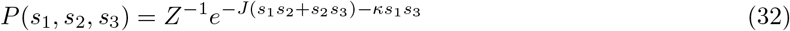

where *Z*^−1^ is the normalization factor. The two-point marginal probabilities are given by

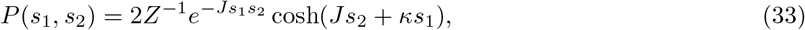

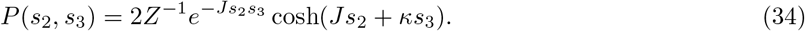

The one-point marginal probability for the middle unit is

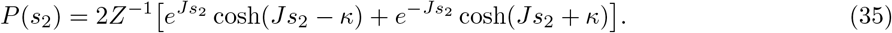

These four probabilities is all we need to quantify the accuracy of the pair approximation, which in our case is represented by *P* (*s*_1_, *s*_2_, *s*_3_) ≈ *P* (*s*_1_, *s*_2_)*P* (*s*_2_, *s*_3_)*/P* (*s*_2_). The numerical accuracy of this approximation can be assessed by defining the ratio *R*_3_ as

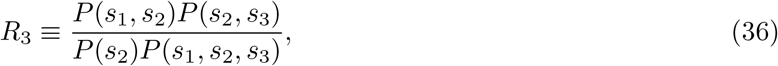

and looking how much *R*_3_ deviates from unity. Since *R*_3_ depends on configurations *s*_1_, *s*_2_, *s*_3_, it is good to determine the mean value of *R*_3_, averaged over all these states, i.e., 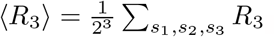.

After some straightforward algebra, we can find ⟨*R*_3_⟩ as

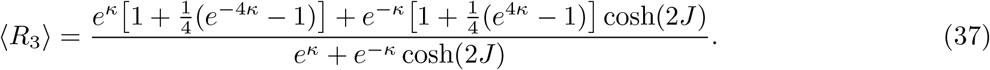

From this formula it is clear that for *κ* ↦ 0 we get ⟨*R*_3_⟩ ↦ 1, regardless of the value of *J*. For large coupling *J* (*J* ≫ 1), we obtain 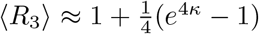, which means that ⟨*R*_3_⟩ is essentially close to 1 for small *κ*. For example, for *κ* = 0.1 we get ⟨*R*_3_⟩ = 1.12, for *κ* = 0.3 we get ⟨*R*_3_⟩ = 1.58, and higher values of *κ* increase ⟨*R*_3_⟩ even further, which breaks the pair approximation. For intermediate values of *J*, e.g. *J* = 1, the ratio ⟨*R*_3_⟩ achieves value 1.55 for *κ* = 0.4, which is a slightly larger value than for the strong coupling case.

## Supplementary Information

The code for performed computations is provided in the Supplementary Material.

## Acknowledgments

The work was supported by the Polish National Science Centre (NCN) grant number 2021/41/B/ST3/04300 (JK).

